# Quaternary structure independent folding of voltage-gated ion channel pore domain subunits

**DOI:** 10.1101/2021.08.15.456357

**Authors:** Cristina Arrigoni, Marco Lolicato, David Shaya, Ahmed Rohaim, Felix Findeisen, Claire M. Colleran, Pawel Dominik, Sangwoo S. Kim, Jonathan Schuermann, Anthony A. Kossiakoff, Daniel L. Minor

**Affiliations:** Cardiovascular Research Institute, University of California, San Francisco, California 93858-2330 USA; Department of Biochemistry and Molecular Biology, University of Chicago, Chicago, IL 60637; Northeastern Collaborative Access Team, Department of Chemistry and Chemical Biology, Cornell University, Ithaca, NY 14850; Departments of Biochemistry and Biophysics, and Cellular and Molecular Pharmacology, University of California, San Francisco, California 93858-2330 USA; California Institute for Quantitative Biomedical Research, University of California, San Francisco, California 93858-2330 USA; Kavli Institute for Fundamental Neuroscience, University of California, San Francisco, California 93858-2330 USA; Molecular Biophysics and Integrated Bio-imaging Division, Lawrence Berkeley National Laboratory, Berkeley, CA 94720 USA

## Abstract

Every voltage-gated ion channel (VGIC) superfamily member has an ion conducting pore consisting of four pore domain (PD) subunits that are each built from a common plan comprising an antiparallel transmembrane helix pair, a short, obliquely positioned helix (the pore helix), and selectivity filter. The extent to which this structure, the VGIC-PD fold, relies on the extensive quaternary interactions observed in PD assemblies is unclear. Here, we present crystal structures of three bacterial voltage-gated sodium channel (BacNav) pores that adopt a surprising set of non-canonical quaternary structures and yet maintain the native tertiary structure of the PD monomer. This context-independent structural robustness demonstrates that the VGIC-PD fold, the fundamental VGIC structural building block, can adopt its native-like tertiary fold independent of native quaternary interactions. In line with this observation, we find that the VGIC-PD fold is not only present throughout the VGIC superfamily and other channel classes but has homologs in diverse transmembrane and soluble proteins. Characterization of the structures of two synthetic Fabs (sFabs) that recognize the VGIC-PD fold shows that such sFabs can bind purified full-length channels and indicates that non-canonical quaternary PD assemblies can occur in the context of complete VGICs. Together, our data demonstrate that the VGIC-PD structure can fold independently of higher-order assembly interactions and suggest that full-length VGIC PDs can access previously unknown non-canonical quaternary states. These PD properties have deep implications for understanding how the complex quaternary architectures of VGIC superfamily members are achieved and point to possible evolutionary origins of this fundamental VGIC structural element.

## Introduction

The voltage gated ion channel (VGIC) superfamily is the largest family of ion channels and encompasses channels that respond to diverse gating cues and that are capable of conducting various cation types^1^. The transmembrane architecture of channels in this superfamily is built from two basic elements. Each channel subunit always contains a pore domain (PD), comprising two transmembrane helices bridged by the pore helix and selectivity filter^2^. Some members also have a voltage-sensor or voltage-sensor like domain (VSD) attached to the PD^3–5^. Regardless of whether a given VGIC superfamily member assembles from multiple, independent subunits as in potassium^2,6^ and TRP channels^3^, or whether all components are encoded into one large polypeptide, as for voltage-gated calcium (Cavs) and voltage-gated sodium channels (Navs)^4^, the ion conductive pore only comes into being upon assembly of four PDs. Extensive structural studies of diverse VGIC superfamily members show that all share the same basic tetrameric pore architecture^2–6^.

Although VGIC are complex multi-component structures, there is a growing appreciation that there are some principles of modularity in their construction. There are numerous examples demonstrating the ability of the four VSD helices to adopt a native-like structure independently from the PD^7–9^. Further, protein dissection studies, particularly for members of the bacterial sodium channel family (BacNavs)^10–14^, have established the independence of the PD tetramer. Such ‘pore-only’ proteins lacking the VSD can fold, assemble, and form functional, selective ion channels^10–13,15,16^. Structural studies of such ‘pore-only’ proteins have shown the canonical quaternary structure found throughout the VGIC superfamily^11,12,17–20^. Nevertheless, whether there are stable intermediate forms and how the four PD subunits assemble into a functional pore is not well understood^21,22^. Labeling studies of Kv1.3^23,24^ and the archaeal channel KvAP^25^ have suggested that a native-like pore helix topology can develop independently of tetramer formation. Although, molecular dynamics simulations have indicated that the basic PD tertiary fold is stable for short periods (∼700 ns)^23^, a VGIC pore domain has never been observed in any form except for the native tetrameric assembly. Further, the extent to which such a structure might be stable outside of its final quaternary assembly is not known.

Here, we present multiple structures of BacNavs pore-only proteins that demonstrate that the fundamental tertiary architecture of the PD, termed the VGIC-PD fold, is able to form a native-like fold independently of the details of its quaternary structure context. Further, we show that the core VGIC-PD fold structural elements are not only found in numerous VGIC superfamily members and channels, but can be identified in other proteins, including both transmembrane and soluble proteins. These findings indicate that the VGIC-PD fold is built from a fundamentally ancient scaffold comprising a pair of anti-parallel helices bridged by a loop. The capacity of such a structure to fold independently of higher order association is in line with long-standing ideas for a two-step folding mechanism for membrane proteins^26,27^ agrees with cysteine accessibility studies suggesting that Kv channel pore domains adopt a native-like topology during folding^23–25^ and points to a possible route for the evolutionary origins of the tetrameric VGIC pore. This structural independence of the VGIC-PD fold has important implications for VGIC biogenesis, understanding disease mutants, and may have a role in transitions between VGIC functional states.

## Results

### BacNav ‘pore only’ structures reveal non-canonical quaternary pore-domain assemblies

We determined crystal structures of three ‘pore-only’ BacNavs: *Alcanivorax borkumensis* NavAb1p^10^; a calcium-selective mutant of *Silicibacter pomeroyi* NavSp1, CavSp1p^10^; and a chimera, NavAe1/Sp1ctdp, having the pore domain of *Alkalilimnicola ehrlichii* NavAe1^18,28^ and the C-terminal domain (CTD) of NavSp1, at resolutions of 2.85Å, 3.5Å, and 4.19Å, respectively (Table S1). All three formed tetramers comprised of pore domain monomers having the canonical tertiary fold encompassing the S5 and S6 transmembrane helices bridged by the P1 and P2 helices and selectivity filter as found in other BacNav structures^14^ (Fig. 1A). However, to our surprise the pore domains in each were arranged in non-canonical quaternary assemblies in which the selectivity filter faced the periphery rather than the central axis (Fig. 1B-D).

**Fig. 1.**
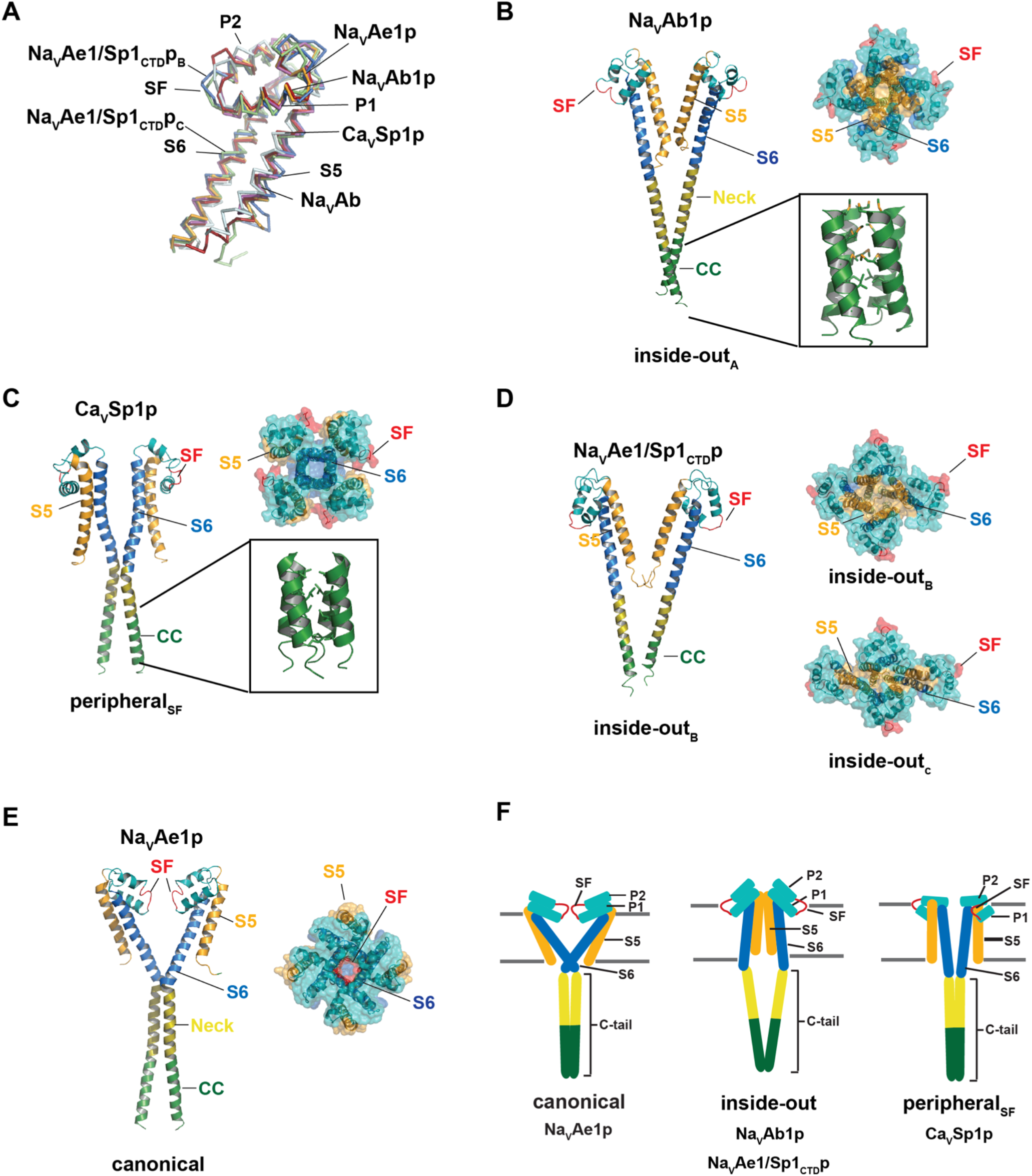
Structures of BacNav PDs. **A,** Superposition of the pore domain subunits from NavAe1p (bright orange) (PDB:5HK7)^18^, NavAb (3RVZ) (magenta)^46^, NavAb1p (firebrick), CavSp1 (pale cyan), NavAe1/Sp1ctdpb (pale green), NavAe1/Sp1ctdpc (marine). **B,** NavAb1p structure, inside-outa form, (left) cartoon showing two of four subunits. Upper right, extracellular view. Inset, coiled-coil. **C,** CavSp1p structure, peripheralsf form, (left) cartoon showing two of four subunits. Upper right, extracellular view. Inset, coiled-coil. **D,** NavAe1/Sp1ctdp structure, inside-outb form, (left) cartoon showing two of four subunits. Upper right and lower left, extracellular view of inside-outb and inside-outc forms, respectively. **E,** NavAe1p structure, canonical form, (PDB:5HK7)^18^ (left) cartoon showing two of four subunits. (Right) extracellular view. F, Cartoon schematics of the canonical, inside-out, and peripheralsf quaternary structures. In B-F channel elements are colored as follows, S5 (bright orange), SF (red), P1 and P2 helices (teal), S6 (marine), neck (olive) coiled-coil (forest).

The NavAb1p structure, termed ‘inside-outa’, showed the most extreme deviation from the canonical pore quaternary structure (Figs. 1B and S1A-C). This quaternary arrangement was observed in NavAb1p crystals grown from detergent that diffracted to 2.85Å (Table S1) and in NavAb1p crystals grown from lipid bicelles that diffracted to 3.64Å (Table S1). The similarity between the detergent and bicelle NavAb1p structures (RMSD_Cα_ = 0.613 and 0.733 for NavAb1p tetramer A and B in bicelles, respectively) (Fig. S1D) and the fact that we could locate a lipid in the 2.85Å structure residing next to a P1 helix aromatic residue where a similar lipid is found in the native pore conformation (Fig. S1E) indicates that the unusual quaternary arrangement was not a consequence of the absence of lipids.

In the inside-outa conformation, each pore-forming domain is rotated ∼180° around the channel central axis relative to the canonical pore domain structure (Figs. 1B and E). This arrangement places the S5 transmembrane helices along the central four-fold axis where they make extensive interactions with each other. Rather than lining the central pore as in the canonical conformation, the S6 helix faces the periphery. These changes result in increased buried surface area of the individual pore domains against their two adjacent neighbors by ∼281 Å^2^ relative to the canonical arrangement found in NavAe1p^11,18^ (Table S2). The neck domain is composed of largely hydrophilic residues and forms a continuous helix that connects the C-terminal end of S6 with the N-terminal end of the coiled-coil. Despite the dramatic rearrangement of pore domain quaternary structure, the NavAb1p coiled-coil domain is folded as in other BacNav structures^11,18^ having the ‘a’ and ‘d’ hydrophobic residues of the coiled-coil sequence^18,29^ in the interior and hydrophilic residues on the exterior (Fig. 1B).

The CavSp1p and NavAe1/Sp1ctdp structures also showed non-canonical quaternary arrangements. CavSp1p crystals were grown from SeMet labeled protein reconstituted in lipid bicelles, diffracted at 3.5Å, and were solved by a two-wavelength MAD experiment (Table S1). CavSp1p shows a quaternary structure in which the transmembrane helices have positions that are similar to those of the canonical pore domain structures, such as NavAe1p^11,18^. S5 is on the periphery and S6 lines the central axis of the tetramer (Figs. 1C and E and S1F-H). However, the pore domain of each subunit has undergone a clockwise rotation of ∼45° relative to the central axis. This change allows the S6 helices to contact each other in a way that closes off central cavity and places the selectivity filter along the exterior of the structure facing bilayer periphery instead of the central pore. Hence, we term this conformation ‘peripheralsf’ (Fig. 1C). The peripheralsf arrangement buries more surface area per monomer (∼123 Å^2^) than the canonical structure (Table S2). There is a kink at Ala226 where the C-terminal end of S6 joins the continuous helix comprising the neck and the coiled-coil domains (Fig. S1I). The CavSp1p coiled coil domain is arranged having its ‘a-d’ hydrophobic repeat in the interior of the four-helix bundle, as seen in the in the NavAe1p CTD^11,18^ (Fig. 1C) and in the NavAb1p ‘inside-out’ forms (Fig. 1B). Hence, as with NavAb1p, the principal quaternary rearrangement from the canonical structure occurs in the pore domain.

NavAe1/Sp1ctdp crystals that diffracted to 4.19Å were solved by MAD using the anomalous signal from selenomethionine containing crystals (Table S1). There were two tetramers in the asymmetric unit that each showed variations of the ‘inside-outa’ arrangement that we term ‘inside-outb’ and ‘inside-outc’ (Figs. 1D and S1J-O). As with the ‘inside-outa’ form, the S5 helix lines the central axis of both ‘inside-outb’ and ‘inside-outc’ NavAe1/Sp1ctdp tetramers while S6 and the selectivity filter face the periphery. The ‘inside-outb’ form is four-fold symmetric and has a wide cavity at the top of the structure, whereas ‘inside-outc’ has a two-fold symmetric arrangement in which the two of the diagonally opposed S5 helices are close to each other while the other two are far apart (Fig. 1D). Contrasting the NavAb1p ‘inside-outa’ and CavSp1p ‘peripheralsf’ forms, the individual pore domain monomers of the ‘inside-outb’ and ‘inside outc’ pore domains bury substantially less surface against their adjacent neighbors relative to the canonical quaternary structure of NavAe1p (-329 and -448 Å^2^, respectively) (Table S2). The neck and coiled-coil regions form helices that are continuous with S6, similar to that observed in the NavAb1p.

Remarkably, the individual pore subunits from this set of four non-canonical quaternary pore domain assemblies formed by three different BacNav pore domains all have tertiary structures similar to those seen in pore domains from canonical quaternary arrangements (Fig. 1A). There are some changes from the native structure in the region of the selectivity filter (Fig. S2). The largest of these encompass residues at the selectivity filter [+1] to [+3] positions in CavSp1p and NavAe1/Sp1ctd. The most obvious change is the dislocation of the [+2] tryptophan. However, this residue is not displaced in NavAb1p (Fig. S2C). These differences suggest that although the overall tertiary structure of the pore domain is independent of the quaternary assembly, attaining the selectivity filter native positions depends on the quaternary structure.

The dramatic differences in the NavAe1/Sp1ctdp pore domain quaternary structures relative to prior NavAe1p structures^11,18^ (Fig. 1D-E) demonstrate that the identical protein sequence, the S5-S6 pore forming element of NavAe1, can adopt both the inside-out and canonical quaternary structure. This preservation of the tertiary fold of the same protein sequence, the NavAe1 pore, in diverse quaternary contexts rules out the possibility that the non-canonical NavAb1p and CavSp1 quaternary structures are peculiar to those particular channels and demonstrates that an individual pore domain can adopt multiple quaternary structures without losing its native tertiary structure. The pore domain monomers in both the NavAe1/Sp1ctd inside-outb and inside-outc conformations bury substantially less surface area against their adjacent neighbors relative to the canonical NavAe1p form (Table S2). This fact rules out the possibility that the preservation of the native-like tertiary structure among the various non-canonical forms results from the non-canonical quaternary contacts. Together, our observations directly demonstrate that the tertiary structure of an individual S5-S6 pore forming element is able to fold into a native-like structure independent of quaternary context.

### VGIC-PD fold structural relatives occur in proteins that are not ion channels

The ability of the BacNav pore monomer to maintain its native-like tertiary structure independent of quaternary context prompted us to ask whether this fold, or something very similar, could be found in other proteins. Hence, we performed a DALI search^30,31^ using the NavAe1p pore monomer (Pro148-Ser285) to find other proteins having clear structural similarity to the S5-S6 PD module. This search identified members of essentially every VGIC superfamily channel for which there is currently structural information, including representatives from the Nav, Cav, and TRP channel families (Fig. 2). This search also identified similarities to channels whose pore domains resemble the VGIC superfamily, such as IP_3_ receptors, glutamate receptors, as well as clearly related ion transport proteins such as the TRKA transporter (Fig. 2). Remarkably, the sequence identity with NavAe1p can be very low (ex. 8% and 11% for GluR and TRPV1, respectively), and yet still yield the same basic architecture comprising two antiparallel transmembrane helices bridged by a loop containing a pore helix (Fig. 2). Notably, in all cases the VGIC-PD monomer occurs in the context of a tetrameric assembly that forms the pore. Due to its predominance in the VGIC superfamily, we denote as the ‘VGIC pore domain fold’ (VGIC-PD).

**Fig. 2.**
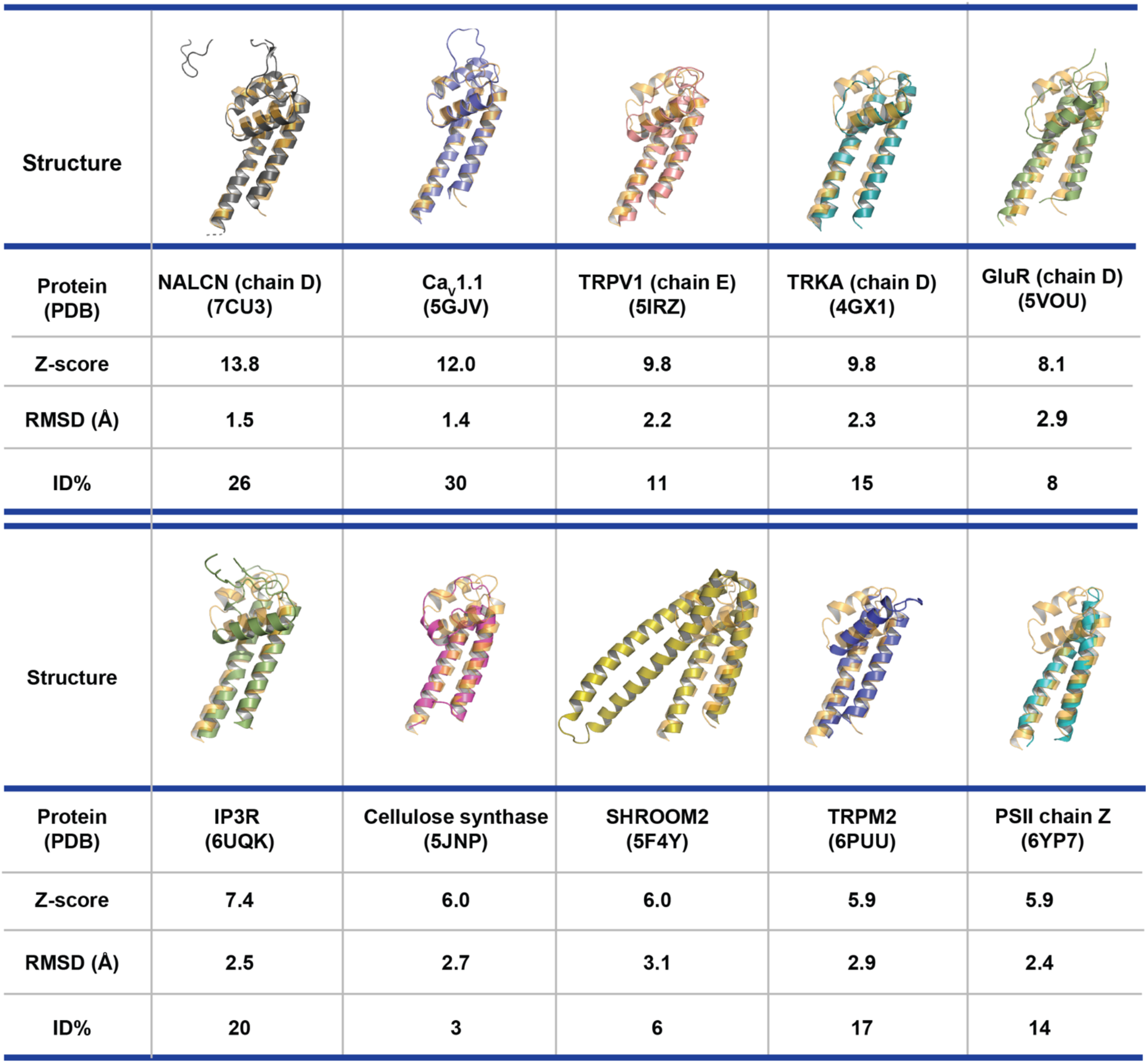
Exemplar structural homologs of the VGIC-PD fold identified using DALI^30,31^. DALI search Z-score, RMSD_Cα_ (Å), and percent identity (ID%) are shown.

To our surprise, the DALI search identified many non-ion channel proteins that had Z-scores and RMSDc*_α_* values that were equivalent to matches between the NavAe1p pore domain and VGIC superfamily members (ex. Z-score ∼6, TRPM2, Fig. 2). The most striking of these was to an element from the plant-conserved region (PC-R) of the rice (*Oryza sativa*) CesA8 cellulose synthase^32^ (Fig. 2). This soluble domain consists of a pair of antiparallel helices bridged by a twenty-five residue loop that contains an *α*-helix and shares structural similarity with NavAe1p that is on par with that of ion channels (RMSDc*_α_* ∼2.5-2.9 Å, Fig. 2) even though the sequence identity with NavAe1p is essentially undetectable (3%). Strikingly, the PC-R CesA8 structure report had noted that high similarity of the interactions made between the connecting loop and the antiparallel helical pair to sodium channels^32^. Other good structural matches from non-ion channel proteins can be seen with the cytoskeletal protein SHROOM2^33^ in which the antiparallel helix pair that matches NavAe1p is bridged by a loop forming a pair of antiparallel helices (Fig. 2) and in membrane proteins such as chain Z of pea (*Pisum sativum*) photosystem II (PSII)^34^ in which the helical pair bridged by a short loop. Together, these observations indicate that a key feature of the structural independence of the VGIC-PD fold tertiary structure is built on the association of the transmembrane antiparallel helical pair. This notion supports earlier ideas regarding the biogenic steps involved in Kv channel pore folding that noted the prevalence of this core feature in both potassium channel pore domains and coiled-coil containing proteins^23^.

### The S6-Neck hinge allows for pore domain rearrangements

The diversity of observed quaternary assemblies prompted us to investigate how these unusual assemblies were related to each other and to the canonical tetrameric pore. Superposition of the cytoplasmic domain of pore domain monomers from NavAb1p, CavSp1p, NavAe1/Sp1ctdp, and NavAe1p^18^ reveals that the different positions of the pore-forming S5-S6 unit are interrelated by simple rigid body movements around a hinge position at the S6-Neck domain junction (NavAb1p His237, CavSp1p Ala223, NavAe1p His245)(Fig. 3A). The canonical arrangement of the individual pore domain subunits in NavAe1p and the CavSp1p pore inside-outsf conformation are connected by a rotation of ∼45° parallel to the plane of the membrane around this point (Fig. 3A-B). An additional tilt of the pore domain by ∼60° parallel to the membrane plane is sufficient to move between the NavAb1p inside-outa conformation and CavSp1p pore inside-outsf (Fig. 3A-B). The NavAe1/Sp1ctdp inside-outb and inside-outc conformations are related to the canonical NavAe1p position in a similar way, involving the 45° rotation parallel to the plane of the membrane and a tilt that places them at intermediate positions between the canonical and inside-outa positions (Fig. S3A-B). Hence, two simple rigid body movements around the hinge at the top of the CTD connect the canonical NavAe1p position to the family of non-canonical pore domain arrangements we observe.

**Fig. 3.**
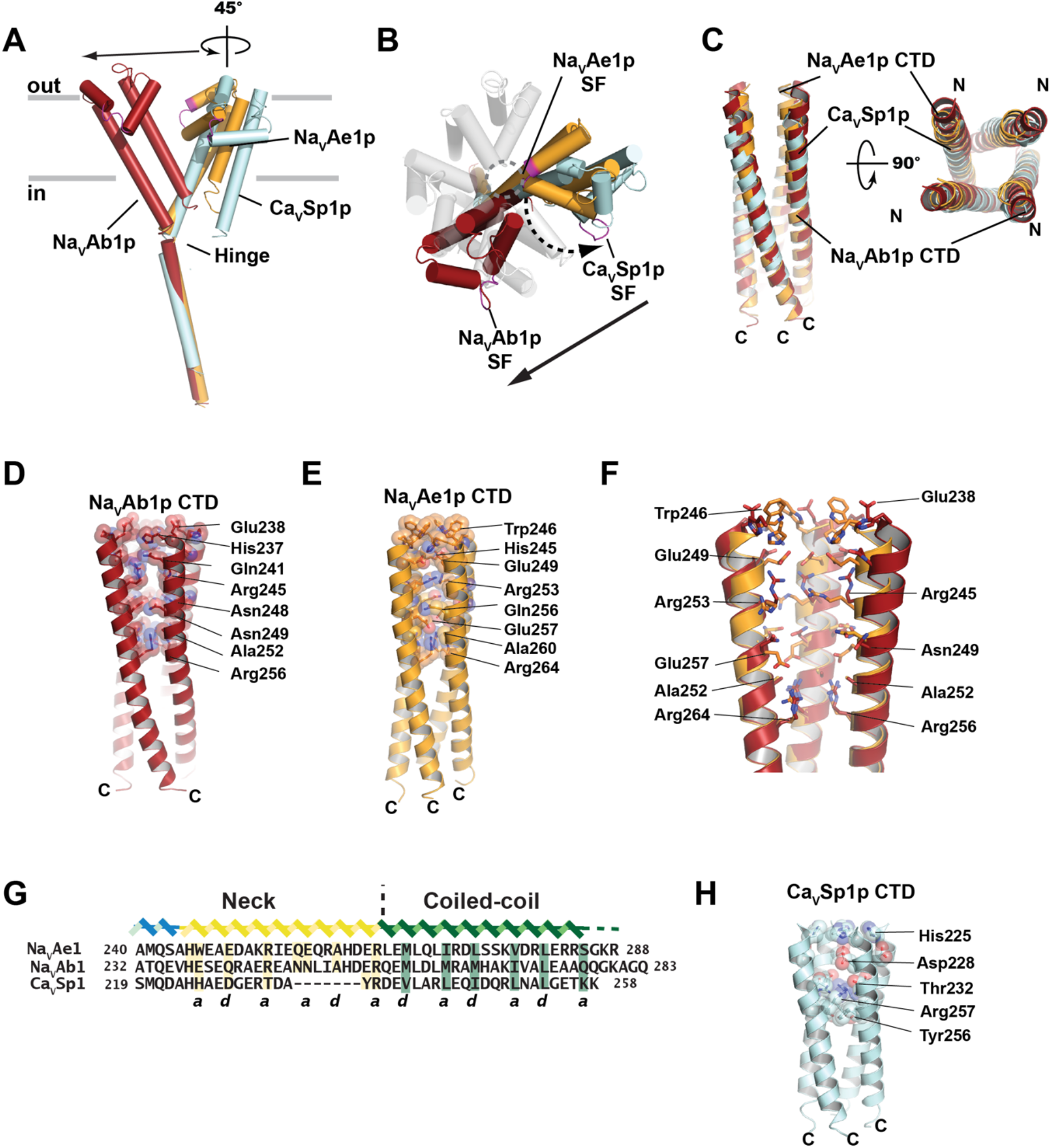
BacNav PD structural relationships and CTD comparisons. **A,** Superpositon showing the rigid body movements that connect NavAe1p (orange), CavSp1p (cyan), and NavAb1p (firebrick) conformations. C-tails of each monomer are superposed. The selectivity filter of each monomer is magenta. Rotation symbol indicates the NavAe1p-CavSp1p relationship. Arrow shows the CavSp1p-NavAb1p relationship. Hinge is indicated. **B,** Extracellular view of ‘A’. Location of central ion conducting pore is indicated by the open circle. Dashed line indicates the NavAe1p-NavSp1p relationship. Arrow shows the NavSp1p-NavAb1p relationship**. C,** Superposition of the CTDs of NavAe1p (orange), CavSp1p (cyan), and NavAb1p (firebrick). **D,** and **E,** CTDs of **D,** NavAb1p and **E,** NavAe1p (orange) showing the neck region hydrophilic cores as space filling. **F,** Detailed view of the superposition of theNavAb1p (firebrick) and NavAe1p (orange) neck region hydrophilic cores. **G,** Sequence alignment of the NavAe1p, NavAb1p, and CavSp1p CTDs. Heptad repeat ‘a’ and ‘d’ positions are indicated. **H,** CavSp1p CTD showing the neck region hydrophilic core in space filling.

Even though there are extreme changes in the quaternary assembly of the pore domains among these structures their CTDs superpose on each other very well (Fig. 3C and S3C). Indeed, analysis of the neck domain regions shows that cores of both NavAb1p and CavSp1p comprise hydrophilic residues having an ‘a/d’ repeat that match the corresponding positions of NavAe1p^18^ (Fig. 3D-G). The main differences from NavAe1p are that the N-terminal end of the NavAb1p neck near the junction with S6 is slightly more open (Fig. 3F) and that the CavSp1p neck is shorter than NavAb1p and NavAe1p by one helical turn (Fig. 3G, 3H, and S3D). The consistent packing patterns of the Neck domain across the structures that show different PD quaternary arrangements, underscore the point that the non-canonical arrangements are facilitated by rigid body motions of VGIC pore domain fold around the hinge at the S6/Neck junction.

### Canonical CTD structures support diverse quaternary assemblies

To understand the differences in quaternary packing better, we used a distance-based analysis to characterize the proximities of various parts of the subunits relative to their neighbors. These contact plots show key intersubunit interactions within the canonical quaternary structure, including extensive interactions between the P1-S6 interface, SF-SF, and the cytoplasmic end of S6 where these helices come together to form the intracellular gate^11,12,35^ (Fig. 4A). None of the non-canonical forms show the P1-S6 or SF-SF contacts present in the canonical assembly (Fig. 4B-F). Taken together with the preservation of the tertiary structure of the VGIC pore domain fold regardless of the quaternary context (Fig. 1A), this analysis highlights that the VGIC pore domain fold can adopt its native structure independent of intersubunit interactions.

**Fig. 4.**
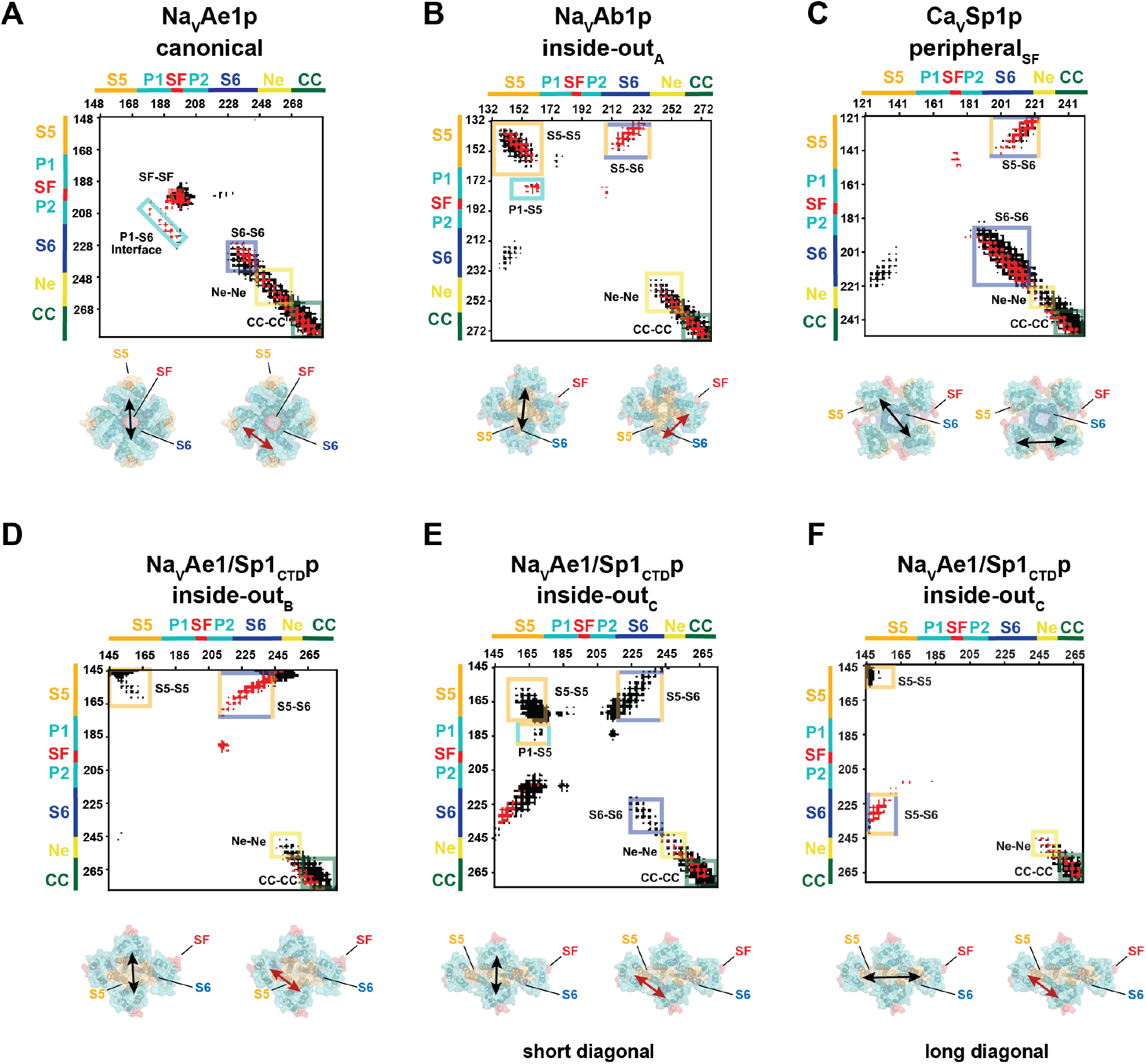
BacNav PD Contact maps. C_α_-C_α_ distances for (black) diagonal subunits at 20 Å and (red) neighboring subunits at 12 Å for **A,** NavAe1p, canonical conformation, **B,** NavAb1p, inside-outa conformation, **C,** CavSp1p, peripheralsf conformation, **D,** NavAe1/Sp1ctdp, inside-outb conformation, **E,** NavAe1/Sp1ctdp, inside-outc conformation for the subunits related by the short diagonal, and **F,** NavAe1/Sp1ctdp, inside-outc conformation for the subunits related by the long diagonal. Channel structural elements are indicated. Extracellular views of the PDs are shown underneath each plot. Arrows indicate the diagonal (black) and neighbor (red) distance relations of the contact plots.

Besides emphasizing the structural independence of the VGIC pore domain fold, the intersubunit distance plots also highlight shared quaternary features among some of the inside-out variants, including close contacts between S5 transmembrane helices in NavAb1p inside-outa (Fig. 4B), NavAe1/Sp1ctdp inside-outc (Fig. 4E) and to a lesser extent in NavAe1/Sp1ctdp inside-outb (Fig. 4D), where the S5 helices are separated at the extracellular end. This feature is absent from the CavSp1p peripheralsf form (Fig. 4C), where as noted above, the S6 segments make extensive contacts that are not present in the canonical form (Fig. 4A). The other key difference from the canonical form is that all of the non-canonical forms show close approaches of S5 and S6 to the neighboring subunit and to the diagonally placed subunit (cf. Fig. 4A and B-F). Contrasting these differences, this analysis highlights the similarities of the neck and coiled-coil regions in all of the ‘pore-only’ BacNavs quaternary forms (Fig. 4). The constancy of the cytoplasmic domain interdomain interactions throughout the canonical and non-canonical structures emphasizes that the non-canonical quaternary arrangements occur due to changes around the hinge between the neck and S6.

### sFabs recognize the VGIC-monomer fold

In an effort to develop anti-BacNav Fabs, we raised synthetic Fabs (sFab) using phage display^8,36^ against purified NavAe1/Sp1ctdp and NavAe1p incorporated into lipid nanodiscs to present the channels in a membrane environment^37^. These efforts yielded sFabs SAT09 and ANT05 from phage display selections using NavAe1/Sp1ctdp and NavAe1p, respectively, and yielded co-crystal structures of complexes with NavAe1/Sp1ctdp (Fig. S4 A-C, Table S1). Crystals of the sFab SAT09:NavAe1/Sp1ctdp diffracted X-rays to 3.6Å. Structure solution by molecular replacement revealed that SAT09 recognizes the selectivity filter and surrounding P1 and P2 helices of the VGIC-PD fold (Fig. 5A) in the context of a tertiary structure for an individual pore domain subunit that is essentially like that found in the canonical NaAe1p structure (RMSDc*_α_* = 2.81 Å vs. NavAe1p (PDB: 5HK7)^18^)(Fig. S5A). The structure shows that residues from both sFab heavy and light chains make extensive contacts with the selectivity filter and P2 helix (Fig. 5B) forming an interface of ∼761Å^2^. The center of these interactions is the conserved selectivity filter [+2] position, Trp199, which, along with the two subsequent residues of the selectivity filter, Ser200 and Met201, is incorporated into the P2 helix (Fig. 5B). This exposed Trp199 conformation is different from its position in the canonical tetrameric pore NavAe1p (Fig. 5C) where it forms an anchor for the native selectivity filter structure and is buried in the intersubunit interface (Fig. 5D) but is similar to what is observed in the inside-out NavAe1/Sp1ctdp structures (Fig. S5B). Hence, the data indicate that this conformation is not induced by the sFab but appears to be a consequence of the sFab recognizing a conformation that is accessible in the absence of canonical intersubunit interactions.

**Fig. 5.**
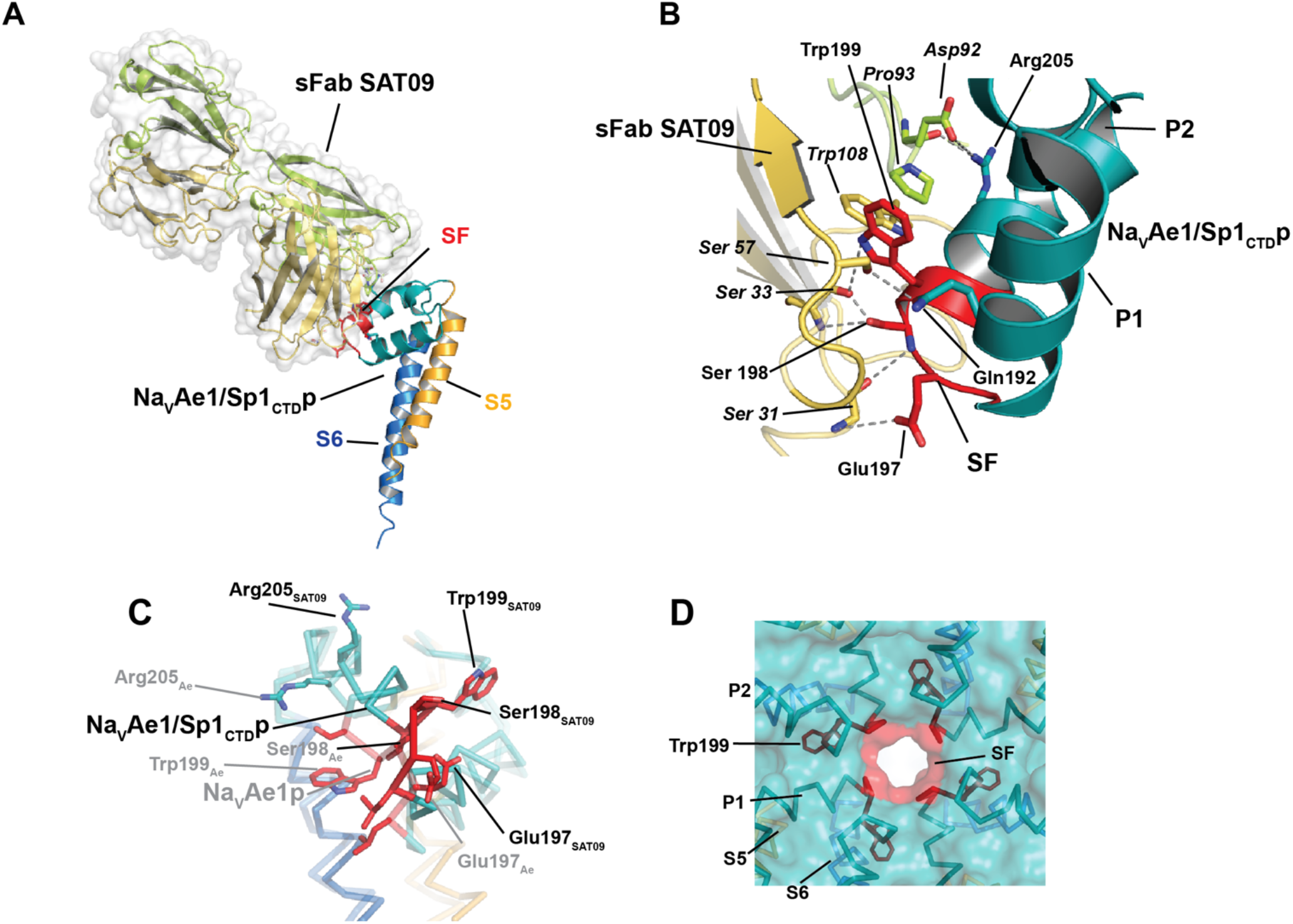
sFab SAT09 recognizes the BacNav SF. **A,** Crystal structure of the sFab SAT09:NavAe1/Sp1ctdp complex. sFab SAT09 is shown with a semitransparent surface, light (limon) and heavy (yellow orange) chains shown as cartoons. NavAe1/Sp1ctdp channel elements are colored as follows, S5 (bright orange), SF (red), P1 and P2 helices (teal), S6 (marine). **B,** Details of the sFab SAT09:NavAe1/Sp1ctdp interface. Colors are as in ‘A’. **C,** Superposition of NavAe1/Sp1ctdp from the sFab SAT09:NavAe1/Sp1ctdp complex and NavAe1p from the canonical structure (PDB:5HK7)^18^. **D,** Quaternary context of sFab SAT09 binding epitope in the NavAe1p canonical structure (PDB:5HK7)^18^.

The sFab SAT09:NavAe1/Sp1ctdp complex formed a decameric structure in the asymmetric unit comprising two head-to-head pentamers held together by extensive sFab SAT09-sFab SAT09 interactions (Fig. S5C). In this arrangement, individual pore domains make very few interactions to each other and are oriented such that the selectivity filter faces the periphery of the complex (Fig. S5D and E). As this pentameric assembly is different from the other non-native forms, we term this arrangement ‘inside-out_D_’ (Fig. S5E). The S6 helix faces the central axis, but comparison with a monomer from the canonical NavAe1p^18^ shows that the NavAe1/Sp1ctdp monomers are tilted by ∼60° around the His245 hinge point between S6 and the neck domain (Fig. S5F and G). There is also a ∼15°twist around the S6 axis. Similar to the other non-canonical NavAe1/Sp1ctdp quaternary arrangements, the individual pore monomers in the inside-out_D_ form bury substantially less surface area against their adjacent neighbors (Table S2), further supporting the idea that the native-like tertiary fold forms essentially independently of the quaternary structure. Contact analysis shows that, in contrast to the other forms, the pore domains make very few interactions, indicating that this quaternary arrangement is largely held together by interactions of the neck domain (Fig. S5H). Notably, the coiled-coil region was not visible, suggesting that the crystallographic pentameric assembly is incompatible with its structure. SEC-MALS analysis (Fig. S6A & B, Table S3) of NavAe1/Sp1ctdp and the SAT09:NavAe1/Sp1ctdp complex shows that both are monodisperse tetramers, indicating that SAT09 does not affect oligomerization state. Hence, these data, together with the low degree of intersubunit contacts in the pentameric assembly suggest that the pentameric form is a consequence of the crystallization conditions and not the result of SAT09 binding to the ‘inside-out’ form.

The low resolution structure of the ANT05:NavAe1/Sp1ctdp complex revealed a similar recognition mode in which the sFab recognizes the selectivity filter and surrounding structure (Fig. S7A). ANT05 binds to an inside-out tetramer and makes substantial interactions with the exposed SF and P1 and P2 helices. Although the resolution of the data preclude detailed description of the interactions, it is clear that ANT05 is rotated ∼90° along the sFab long axis relative to SAT09 (Fig. S7B). Hence, this represents a different recognition mode of the same structural epitope targeted by SAT09, consistent with the sequence differences between the complementary determining regions (CDRs) of sFabs SAT09 and ANT05 (Fig. S4C). Notably, ANT05 was generated from selections using NavAe1p, a ‘pore-only’ channel for which we have never observed non-canonical quaternary structures, even when the neck region structure was disrupted and rendered flexible^18^. The fact that we were able to raise an sFab, ANT05, that recognizes a non-native quaternary structure using the natively assembled NavAe1p^18^ as the selection target suggests that even natively-assembled pores can access non-canonical quaternary forms.

### sFab SAT09 recognizes full-length BacNavs in multiple contexts

Given our repeated observations of non-native quaternary structures in different BacNav ‘pore-only’ constructs, we wanted to probe whether such conformations could exist in the context of a full-length channel. The selectivity filter [+2] Trp that is at the center of the epitope recognized by SAT09 is inaccessible in the canonical quaternary structure due to its burial in the intersubunit interface (Fig. 5D). Superposition of the SAT09:NavAe1/Sp1ctdp complex onto the canonical NavAe1p pore reveals that even if the [+2] Trp could flip out to be exposed in the canonical conformation, SAT09 would make extensive clashes with the surrounding subunits that would prevent this sFab from binding to a BacNav pore when assembled in the canonical tetrameric structure (Fig. S8A). Thus, we took advantage of the ability of SAT09 to bind to the exposed SF epitope to test whether this sFab could bind to full-length BacNavs.

We first used Fluorescence-detection Size Exclusion Chromatography (FSEC)^38^ to test whether SAT09 could bind to detergent-solubilized C-terminal GFP fusions of three purified full-length BacNavs: NavAe1, NavSp1, and NavAe1/Sp1ctd (Fig. 6A). For all three full-length channels, inclusion of SAT09 shifted the elution volume in a manner consistent with the formation of a complex (Fig. 6B). Importantly, we were not able to detect such a shift using K_2P_2.1 (TREK-1)^39^, a well-characterized, folded, purified potassium channel that lacks the SAT09 binding epitope, as a control (Fig. S8B). Hence, these data indicate SAT09 is able to recognize its SF epitope in the context of a full-length channel and suggest that non-canonical pore conformation can be accessed in the presence of the voltage sensor domain.

**Fig. 6.**
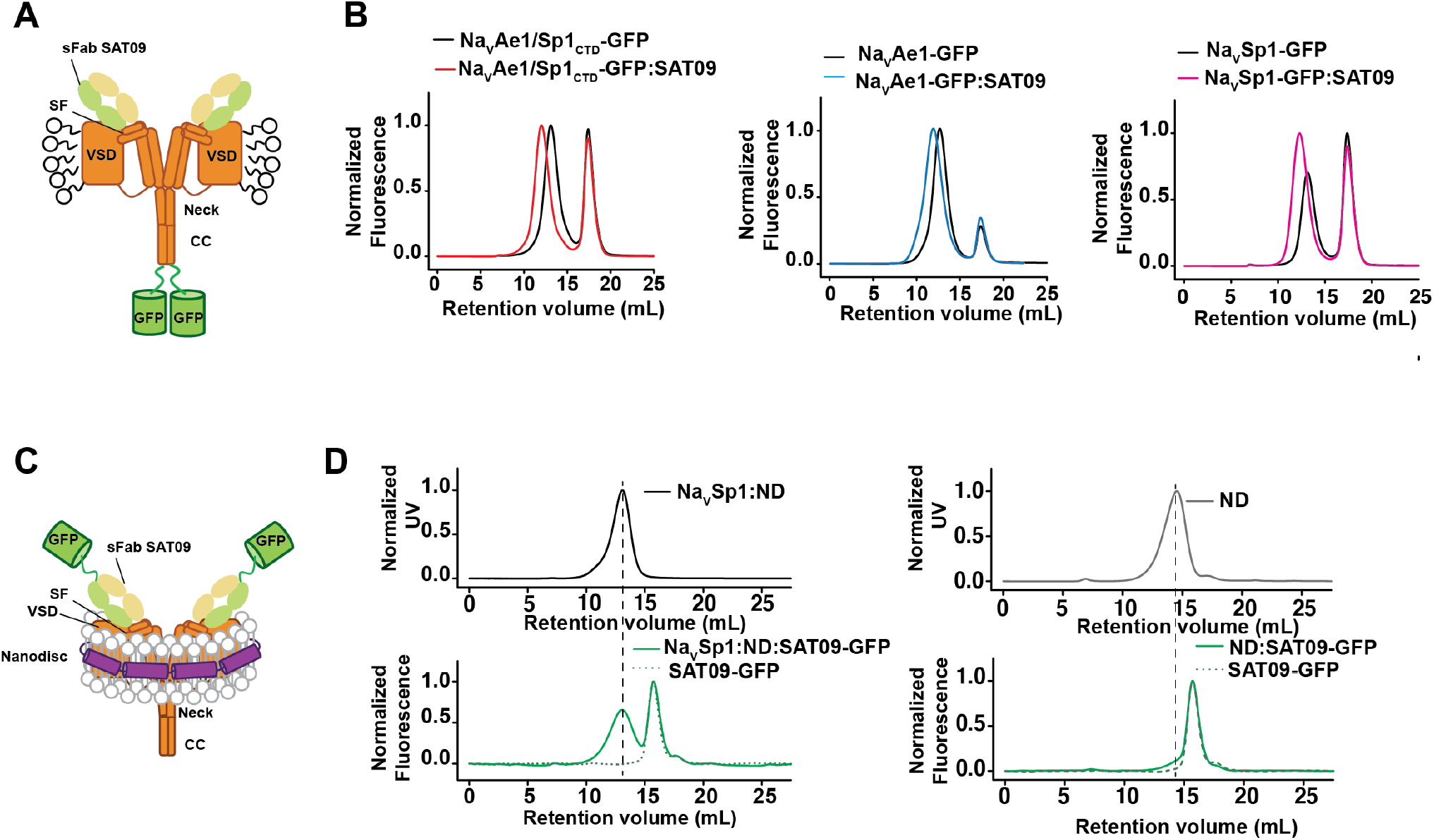
Evidence for non-canonical pore domain quaternary structure in full-length BacNavs. **A,** Cartoon showing interaction of sFab SAT09 with an ‘inside-out’ pore domain conformation in the context of a full-length BacNav in a detergent micelle. **B,** Superose 6 FSEC profiles for GFP-tagged full-length channels NavAe1/Sp1, NavAe1, and NavSp1 alone and with 10 µM sFabSAT09. **C,** Cartoon showing interaction of sFab SAT09 with an ‘inside-out’ pore domain conformation in the context of a full-length BacNav in a lipid nanodisc. **D,** Superose 6 profiles monitored by (top) UV absorbance at 280 nm for NavSp1:nanodiscs (ND) and ND alone and (bottom) FSEC for 10 µM sFabSAT09 alone (green dashed line) and with NavSp1:ND or ND alone.

We next tested the ability of SAT09 to recognize full-length BacNavs in a lipid environment. Of the three channels that SAT09 could recognize in detergent, only NavSp1 could be readily incorporated into soybean lipid nanodiscs. Because the large size of the NavSp1:ND complex would make it difficult to observe a substantial shift in elution volume upon addition of the 50kDa sFab, we adopted a different strategy to detect the interaction. We used the Spycatcher strategy^40^ to make a covalent attachment between purified GFP bearing the SpyCatcher domain and an sFab SAT09 construct bearing the SpyTag peptide at its N-terminus. We then used a combination of UV absorbance and FSEC to evaluate whether the purified GFP-tagged SAT09 would recognize NavSp1 reconstituted into lipid nanodiscs (Fig. 6C). Comparisons of the UV absorbance profiles that show the migration of NavSp1:nanodisc complex with the FSEC profiles run in the presence of SAT09-GFP show that SAT09 co-elutes with the nanodisc containing-fraction only when NavSp1 is present (Fig. 6D), indicating specific recognition of the channel in a lipid environment. To investigate the stoichiometry of the SAT09:NavSp1 interaction, we used SEC-MALS analysis. Because of difficulties analyzing nanodiscs using this method, we replaced the nanodics with the amphipol A8-35^41^. These data reveal two monodisperse peaks for NavSp1, one that had molecular mass matching a tetrameric channel and a higher molecular weight peak that was consistent with a dimer of channel tetramers (Fig. S7C and Table S3). Notably, the addition of SAT09 shifted both peaks to higher molecular weight species, with the peak containing the channel tetramer displaying a mass that was consistent with a stoichiometry of 2 SAT09 sFabs to one channel (Table S3). These data indicate that in the context of a full-length channel, it is possible to have at least two pore domain subunits in which the inside-out-like form is present. Because binding the epitope of SAT09 is incompatible with the native quaternary form (Fig.S8A), taken together, these results indicate that inside-out like conformations can be accessed by some subunits in the context of the full-length channel.

## Discussion

Our studies of the structures of ‘pore-only’ BacNavs yielded a surprising array of well-folded, protein assemblies having unconventional PD quaternary structures. These structures were a surprise as multiple functional studies of ‘pore-only’ constructs, including those having non-canonical structures determined here, NavAbp1 and CavSp1p, form functional, selective ion channels^10–13,15,16^. The striking preservation of the native-like PD monomer structure in these diverse assemblies in which varied types of intersubunit contacts are made support the idea this fold, termed the VGIC-PD fold, is able to adopt its native tertiary structure in the absence of specific quaternary arrangement. These structural observations are in agreement with prior inferences about the potential stability of the native-like structure of the Kv1.3 PD made from a combination of labeling studies and simulation^23,24^. The stability of the PD tertiary fold also aligns the idea that the structures of many transmembrane domains are under thermodynamic control^42,43^ and is consistent with ideas from the two-stage model of membrane protein folding in which helical pairs are able to form and associate prior to assembly into the final native structure of the complete transmembrane protein^26,27^.

All VGICs require four PDs to make a functioning ion conducting pore assembled from either multiple, independent polypeptide chains, as in the case of voltage-gated potassium channels^2^ and TRP channels^3^, or from chains that bear multiple PD copies, such as K_2P_^6^, TPC^5^, and voltage-gated calcium and sodium channels^4^. This requirement means that regardless of whether the PDs come from one or multiple polypeptides, there is some period of time in which the PDs must exist following their initial synthesis and membrane insertion without their final partners that form the tetrameric channel pore. The average rate of eukaryotic protein synthesis rate is ∼5.6 aa/sec^44,45^. If one considers a channel having all four pore domains within a single polypeptide, such as the voltage-gated sodium channel Nav1.4, the time between the completion of synthesis of the first PD (PD1) and fourth PD (PD4)(spanning ∼1140 aa) is ∼3.4 minutes. The time between completion of PD1 and the C-terminal ends of the other PDs is also substantial, being ∼1 min to end of PD2 and ∼2.8 min to end of PD3. These constraints are unlikely to have major consequences on the VSDs as their self-contained four-helix bundle structure channel can fold independently^7–9^. However, the consequence for the PDs is more serious, as three of the four PDs would have to wait many minutes until they could adopt their final folded quaternary structure upon completion of the synthesis of PD4. The ability of the VGIC-PD fold to adopt its native structure independent of its final native quaternary context is likely to be important for the process of the maturation of the final native state of VGICs, and may help to protect these domains from being recognized as damaged or misfolded proteins that need to be eliminated from the membrane.

The observation that the SAT09 sFab is able to bind NavSp1 indicates that non-canonical pore arrangements can be accessed in the context of a full-length channel and raises the possibility that such conformations could impact function. Prior electron paramagnetic resonance (EPR) studies of NavSp1 and NavSp1p noted substantial mobility differences between the pore domains of NavSp1 and NavSp1p^15^. Residues reported to show the largest mobility changes as well as enhanced lipid accessibility include three P1 pore helix residues that are buried in the canonical structure intersubunit interface but that are exposed in the non-canonical conformations, including the ‘peripheralsf’ CavSp1p conformation (Fig. S9). Supported by the observation that the F170C P1 helix mutation enhanced slow inactivation, Chatterjee *et al*. interpreted the differences in dynamics and lipid accessibility between NavSp1 and NavSp1p as indicative of a slow inactivated conformation in NavSp1p^15^. Based on their exposure in the CavSp1p structure (Fig. S9), it seems likely that the EPR parameter differences originate from non-canonical conformations similar to those we observe. Whether such conformations are slow inactivated states remains unclear. The scale of these changes contrasts with the smaller, largely rigid body conformational changes in the pore domains of BacNavs that have also been suggested to represent an inactivated channel^46–48^. However, it is worth noting that conformational changes from the canonical to non-canonical quaternary structures only require rigid body movements (Fig. 3A and B). This property together with the observation that sFab SAT09 binding indicates that at least two pore domain subunits can access a non-canonical arrangement in a full-length channel. Taken together with the pore domain mobility results of Chatterjee *et al*^15^, these observations raise the possibility that the manifold of conformational states that underlies slow inactivation may be richer than previously thought.

Our observation of the robust, quaternary structure-independent nature of the VGIC-PD tertiary fold sets a new framework for thinking about the fundamental structural units that comprise VGICs, their biogenesis, and evolutionary origins. Our data show that, similar to the other major functional unit in the VGIC transmembrane architecture, the VSD^7–9^, the VGIC-PD tertiary fold can adopt a native-like structure without the requirement to be assembled into its final, functional tetrameric form. This independence may have relevance for the steps required to assemble a functioning VGIC from its constituent parts. Further, disease-causing channel mutations may exert their effects by disrupting this structure or its ability to partner with other subunits. Perhaps the most profound insight from the demonstration of the robustness of the VGIC-PD fold is the consequence of this property for the origins of the VGIC superfamily. The existence of structurally similar units in soluble enzymes as well as other proteins that are not channels (Fig.2) suggests that oligomerization, followed by some optimization of the interhelix loop were the key steps in the origin of the VGIC tetramer. The observations we describe here should serve as the starting point for new studies of ion channel biogenesis, evolution, and the origins of still poorly understood functional states.

## Acknowledgements

We thank K.Brejc, L. Jan, and M. Grabe for comments on the manuscript. S. Wong for expert technical assistance in protein preparation. This work was supported by grants NIH-NHLBI R01-HL080050, NIH-NIDCD R01-DC007664, and the Program for Breakthrough Biomedical Research, which is partially funded by the Sandler Foundation to D.L.M., NIH-NIGMS GM117372 to T.K., and an AHA postdoctoral fellowships to C.A and M.L.

This work is based on research conducted at the Northeastern Collaborative Access Team beamlines, which are funded by the National Institute of General Medical Sciences from the National Institutes of Health (Grant P30 GM124165). This research used resources of the Advanced Photon Source, a U.S. Department of Energy (DOE) Office of Science User Facility operated for the DOE Office of Science by Argonne National Laboratory under Contract DE-AC02-06CH11357.

## Author Contributions

C.A. and D.L.M. conceived the study and designed the experiments. D.S. purified and crystallized the initial structures of CavSp1p and NavAb1p in detergent. A.R. purified, crystallized, and determined the structure of NavAb1p in bicelles. M.L. determined the structures of the NavAe1_Sp1CTD_p and the SAT09 and ANT05 complexes, and refined all of the structures. F.F. determined structures of CavSp1p and NavAb1p in detergent. C.M.C. and C.A. expressed an purified the proteins and sFab complexes. C.A. crystallized NavAe1/Sp1ctdp and the SAT09 and ANT05 complexes and performed the biochemical characterization. P. D. and A. K. provided the platform for the development of sFabs. P.D. selected the sFabs. J. S. contributed to the ANT05 complex data collection and structure determination. D.L.M analyzed data and provided guidance and support. C.A., M.L., and D.L.M. wrote the paper.

## Data and materials availability

Coordinates and structures factors for NavAb1p (DM), NavAb1p (bicelles), CavSp1p (bicelles), NavAe1/Sp1ctdp (DDM), NavAe1/Sp1ctdp:SAT09, and NavAe1/Sp1ctdp:SAT09 are deposited in the RCSB under accession codes NavAb1p (DM) (PDB:7PGG), NavAb1p (bicelles) (PDB:7PGI), CavSp1p (bicelles) (PDB:7PGF), NavAe1/Sp1ctdp (DDM) (PDB:7PGH), NavAe1/Sp1ctdp:SAT09 complex (PDB: 7PGP), and NavAe1/Sp1ctdp:ANT05 complex (PDB:7PG8) and will be released immediately upon publication.

## Materials and Methods

### DNA constructs

‘Pore-only’ constructs for *Silicibacter pomeroyi* CavSp1p, *Alkalilimnicola ehrlichii* NavAe1p, and *Alcanivorax borkumensis* NavAb1p are as described in^10^. NavSp1 and NavAe1 full-length were cloned from *Silicibacter pomeroyi* and *Alkalilimnicola ehrlichii* genomic DNA, respectively, into pET24b to create a fusion protein having in series, a Tobacco Etch Viral (TEV) Protease site, Green fluorescent protein (GFP), and His_6_ tag at the C-terminal end. The NavAe1_Sp1CTD_ chimera is identical to that reported previously^18^ having the NavAe1 transmembrane region, residues 1 to 241, joined to the NavSp1 CTD, residues 221-258. NavAe1_Sp1CTD_p spans NavAe1_Sp1CTD_ residues 142 to 258 and was cloned in a HM3C-LIC pET24b-derived vector^49^ similar to the other ‘pore-only’ constructs.

### Protein expression and purification

Expression and purification of CavSp1p, NavAe1p, NavAb1p, NavAe1_Sp1CTD_p was carried out as described in^10^. The NavAb1p construct that lead to the highest resolution crystal structure had a double alanine deletion in the coiled-coil (*Δ*Ala275/Ala276) that improved crystal packing. Proteins were expressed in *Escherichia coli* C41(DE3) grown in 2YT (5 g NaCl, 16 g tryptone, 10 g yeast extract) containing 25 µg mL^−1^ kanamycin. Cultures were inoculated with 1 mL L^−1^ of an overnight starter culture grown from a freshly transformed single colony and grown at 37 °C to OD_600 nm_ 0.5, at which point the growth temperature was reduced to 18 °C. After ∼30 minutes at 18 °C, cultures were induced by addition of isopropyl β-D-1-thiogalactopyranoside (IPTG) to 0.4 mM and grown for 52 h. Cells were harvested by centrifugation (6,000 × g at 4 °C). Cell pellets were frozen in liquid N_2_ and stored at −80 °C. Frozen cell pellets were thawed on ice, resuspended in 250 mL cold lysis buffer (300 mM NaCl, 1 mM phenylmethylsulfonyl fluoride (PMSF), 50 mM Tris-HCl pH 8.0,) and disrupted using an EmulsiFlex-C5 homogenizer (Avestin). Cell lysates were cleared from unbroken cells and debris by centrifugation (10,000 × g at 4 °C, 1 h) and the supernatant was then ultracentrifuged (160,000 × g at 4 °C, 2 h) to pellet and separate the membranes from the supernatant, which was discarded. Membranes were homogenized in 50 mL of storage buffer (200 mM NaCl, 8% (vol∕vol) glycerol, 20 mM Tris-HCl pH 8.0) with a Dounce Tissue Grinder (Kimble Kontes LLC), frozen in liquid N_2_ and stored at −80 °C. Frozen membranes were thawed on ice and extracted by addition of detergent in powder form, n-dodecyl-β-D-maltopyranoside (DDM, Anatrace) for NavAe1p and NavAe_Sp1CTD_p and n-decyl-β-D-maltopyranoside (DM, Anatrace) for CavSp1p and NavAb1p, to a final concentration of 20 mM at 4 °C for 2 h with gentle agitation. Detergent-solubilized membrane proteins were separated from insoluble material by ultracentrifugation (160,000 × g at 4 °C, 1 h), and the supernatant was collected.

Detergent extracted His-MBP-tagged proteins were applied to a 40 mL POROS MC 20 Ni^2+^ column (Applied Biosystems) equilibrated with buffer A (200 mM NaCl, 8% (vol∕vol) glycerol, 1.5 mM DDM or 5mM DM, 20 mM Tris-HCl pH 8.0). The column was washed with seven column volumes (CVs) of buffer A supplemented with 50 mM imidazole. The bound proteins were eluted by step application of buffer A supplemented with 300 mM imidazole over two CVs. The affinity tag was removed with an in-house purified His-tag labeled 3C protease^10^ at ratio of 10∶1 (wt∕wt) fusion protein∶protease at 4 °C for overnight with gentle agitation. The following day imidazole was removed using a HiPrep 26/10 Desalting Column (GE Healthcare) preequilibrated with buffer A, and the cleaved protein was separated from the affinity tags and the protease by passing it through the POROS MC 20 Ni^2+^ column (Applied Biosystems) in tandem with a 20-mL amylose column (New England Biolabs) equilibrated in buffer A. The protein was concentrated to 15 mL using an Amicon Ultra-15 centrifugal filtration device (50-kDa MW cutoff, Millipore). The protein was diluted with a modified buffer A (8% (vol∕vol) glycerol, 1.5 mM DDM or 5mM DM, 20 mM Tris-HCl pH 8.0) to a final volume of 50 mL and a final NaCl concentration of 60mM. The sample was loaded onto a 10-mL POROS HQ ion exchange column (Applied Biosystems) equilibrated with buffer B (60 mM NaCl, 8% (vol∕vol) glycerol, 1.5 mM DDM or 5mM DM, 20 mM Tris-HCl pH 8.0). The protein was eluted by a linear gradient from 60 to 500 mM NaCl over 15 column volumes, concentrated using an Amicon Ultra-15 centrifugal filtration device (50-kDa MW cutoff, Millipore) and applied to a Superdex 200 10/300 GL column (GE Healthcare) in buffer C (200 mM NaCl, 0.3 mM DDM or 2.7 mM DM, 20 mM Hepes pH 8.0). Sample purity was evaluated by SDS-PAGE stained with Coomassie brilliant blue R-250. Protein concentration was determined by absorbance at 280 nm using a Nanodrop 2000c spectrophotometer (Thermo Scientific).

### Synthetic Fab (sFab) selection and purification

sFabs were generated from a phage display library, sorting procedure and selection as described previously^50^. Briefly, His_6_-MBP tagged^10^ NavAe1p and NavAe1_Sp1CTD_ were reconstituted in eggPC:POPC (1:4) MSP 1D1^51^ biotinylated nanodiscs. After removal of the empty nanodiscs via Ni-NTA purification, the biotinylation efficiency of nanodiscs was evaluated by pull-down on streptavidin-coated magnetic beads. Reconstituted BacNavs were used for phage library sorting using sFab Library E (kindly provided by S. Koide^52^ based on described protocols^37^).

Single-point phage ELISA was used in the primary validation of binding affinities of generated sFabs in phage format as described previously^37^. Amplified phage particles at 10-fold dilution were assayed against 20 nM biotinylated membrane proteins in nanodiscs using horse radish peroxidase (HRP)-conjugated anti-M13 monoclonal antibody (GE Healthcare, #27-9421-01). Assays were performed in library sorting buffer (200 mM NaCl, 25 mM Hepes, pH 8.0) supplemented with 2% BSA. Each sFab clone with A_450_ signal above 0.2 (three times the average background level of the assay) was sequenced and unique sFabs were sub-cloned into pSVF4 or pIPTG vectors.

Expression and purification of SAT09 and ANT05 H12 sFabs were carried out as described in^53^. The ANT05 H12 variant was derived from ANT05 using a strategy to reduce flexibility in the “elbow” linker region between the Fab variable and constant domains^54^. This variant replaces the heavy chain V^111^SSASTKG^118^ sequence with a sequence, V^111^FN-QIKG^118^, bearing both mutations and a deletion. The dash indicates the position of the deleted residue.

In brief, sFabs were expressed in *E.coli* BL21 Gold grown in 2YT containing 100 ug mL^-1^ ampicillin at 37°C to OD_600nm_ = 0.8 and expression was induced with 1 mM IPTG at 37°C for 4 hours. Cells were harvested, flash-frozen in liquid N_2_ and stored at -80°C.

For sFab purification, pellets were resuspended in 100 mL of phosphate buffer PBS (500 mM NaCl, 20mM Sodium Phosphate, pH 7.4), supplemented with 10 µg mL^-1^ DNase I, 0.5 mM MgCl_2_ and 1 mM PMSF. Cell lysis was performed with a high-pressure homogenizer EmulsiFlex-C5 (Avestin) and the lysate was incubated for 20 minutes at 60°C to precipitate endogenous proteins and Fab degradation products. Lysate was cooled down on ice and ultracentrifugated at 160,000xg for 1hr at 4°C and loaded onto a 5 mL Protein A (GE Healthcare) column equilibrated with PBS. Column was washed with 10 CVs of PBS and protein was eluted with 2 CVs of 100 mM acetic acid. The elution sample was directly loaded onto a 20 mL Poros HS column (Applied Biosystems) packed in house for cation exchange chromatography. The column was washed with 5 CVs of 50mM sodium acetate pH 5 and the Fab was eluted by a linear gradient from 0 to 1 M NaCl. Fractions containing the eluted protein were loaded onto a HiPrep desalting column (GE Healthcare) equilibrated with PBS. Fabs were concentrated to 20 mg mL^-1^, flash-frozen in liquid N_2_ and stored at -80°C.

### Reconstitution of NavSp1 in nanodiscs

NavSp1 expression and membrane preparation was performed as described for BacNavs pores. Frozen membranes were thawed on ice and extracted by addition of solid n-dodecyl-β-D-maltopyranoside (DDM, Anatrace) to a final concentration of 20 mM at 4 °C for 2 h with gentle agitation. Detergent-solubilized membrane proteins were separated from insoluble material by ultracentrifugation (160,000 × g at 4 °C, 1 h), and the supernatant was collected.

Detergent extracted His-MBP-NavSp1 was applied to a 40 mL POROS MC 20 Ni^2+^ column (Applied Biosystems) equilibrated with buffer A (200 mM NaCl, 8% (vol∕vol) glycerol, 1.5 mM DDM, 20 mM Tris-HCl, pH 8.0). The column was washed with seven column volumes (CVs) of buffer A supplemented with 50 mM imidazole. The bound protein was eluted by step application of two CVs of buffer A supplemented with 300 mM imidazole. The eluted protein fractions were concentrated to ∼12 mL using an Amicon Ultra-50 centrifugal filtration device (100-kDa MW cutoff, Millipore) and loaded onto a HiPrep Desalting column 26/10 (GE Healthcare) to remove imidazole. The protein was concentrated to 85 µM and mixed with purified MSP2N2 (expressed and purified as described in^55^), and soybean lipid extract (Avanti Polar Lipids) in a 1:1:150 molar ratio. Following incubation at 4°C for 1 hour, Biobeads (Biorad) that had been pre-washed (30 minutes in methanol, 30 minutes in water, followed by 30 minutes in 200mM NaCl, 20mM Tris-HCl, pH8.0) were added to the mixture and the sample was incubated at 4°C for 1 hour. To ensure nanodisc formation and complete removal of detergent Biobeads were replaced after 1 hour and the sample was incubated overnight. The following day the sample was recovered by carefully pipetting to avoid the Biobeads and loaded onto a 10 mL-amylose gravity column. Column was washed with 10 CV of a buffer containing 200mM NaCl, 20mM Tris-HCl, pH8.0. Purified 3C protease^10^ was added overnight at 4°C to achieve on-column cleavage. The following day the flow-through containing NavSp1 reconstituted in nanodiscs was collected, concentrated to a volume of 500 µL using an Amicon Ultra-50 centrifugal filtration device (100 kDa MW cutoff, Millipore), loaded onto a Superose 6 30/100 gel filtration column and relevant fractions were collected for FSEC assay.

### Generation of GFP-fused SAT09 Fab

We used the SpyTag-SpyCatcher system^40^ to create a GFP-tagged SAT09 Fab for FSEC assays. The SpyTag was inserted at the SAT09 N-terminus just after the periplasm export sequence and the modified SAT09 was expressed as above for untagged SAT09. GFP was fused to the C-termini of the SpyCatcher sequence subcloned in pDEST14 (Addgene plasmid # 35044) and expressed in *E.coli* BL21 (DE3). Briefly, 2L of 2YT media supplemented with 100 µg mL^-1^ ampicillin were inoculated with 20 mL of an overnight starter culture grown from a freshly transformed single colony and incubated at 37°C, 200rpm until OD_600nm_ = 0.5. The incubation temperature was lowered to 30°C and 0.5mM IPTG was added to induce protein expression. After 3 hours, cells were harvested by centrifugation (6000 xg, 4°C, 30min). After sonication debris were removed by ultracentrifugation (160,000 ×g at 4 °C, 1 h), and the supernatant was loaded onto a 40 mL POROS MC 20 Ni^2+^ column (Applied Biosystems) equilibrated with buffer B (300 mM NaCl, 50 mM Tris-HCl, pH 8.0). The column was washed with five CVs of buffer B supplemented with 50 mM imidazole. The bound protein was eluted by step application of buffer B supplemented with 300 mM imidazole over two CVs. 25 µM of purified SpyCatcher-GFP was mixed with excess purified SpyTag-SAT09 (1:1.5) and incubated at 4°C for 1 hr and loaded on a HiPrep Desalting Column (GE Healthcare) equilibrated with 10mM NaCl, 10mM Tris-HCl, pH 8.0 to lower the salt concentration. The complex was purified by ion-exchange on a 4mL-PorosQ equilibrated in 10mM NaCl, 10mM Tris-HCl, pH8.0 and eluted over a gradient of 20 CVs to a final concentration of 500mM NaCl. The sample was loaded on a HiPrep Desalting Column (GE Healthcare) equilibrated with 200mM NaCl, 10mM Tris-HCl, pH 8.0, flash-frozen in liquid N_2,_ and stored at -80°C.

### FSEC and MALS assay

NavSp1, NavAe1 and NavAe1Sp1CTD with a C-terminal GFP tag subcloned in pET24b were transformed in *E.coli* C41 (DE3) for small scale expression to test the binding of SAT09 with fluorescent size exclusion chromatography (FSEC)^38^. Small scale expression followed the same protocol described for the purification of BacNavs pores, with the difference being that only 50mL of culture were inoculated for protein expression. Following sonication cell membranes were isolated by ultracentrifugation (160.000xg, 2hrs, 4°C), resuspended in a solubilization buffer containing 200mM NaCl, 10mM DDM, 20mM Hepes, pH8 and incubated for 2 hours at 4°C. The suspension was ultracentrifuged (160.000xg, 1hr, 4°C) and loaded onto a Superose 6 10/300 connected to a Shimadzu LC-20AD system having an RF-10XL fluorescence detector and SPD-20A UV/Vis detector. Elution profiles of full-length channels were compared with profiles of samples where each channel was incubated with SAT09-GFP overnight at 4°C. As control, the experiment was performed on an unrelated ion channel, K_2P_2.1_cryst_ (TREK-1) that was expressed purified as described previously^39^.

In order to test SAT09-GFP binding on NavSp1 FL reconstituted in nanodisc we observed the shift in molecular weight of the fluorescent Fab co-eluting with the channel confirming the formation of the Fab-channel complex. For each run 1.1 nmol of reconstituted channel was incubated with 1 nmol of Fab. These amounts were empirically determined to ensure good noise-to-signal ratio avoiding saturation of fluorescent signal of unbound Fab. As negative control 2.2 nmoles of empty nanodisc were incubated with 2 nmoles of SAT09-GFP.

### Protein Crystallization and structure determination

#### Crystallography

For all the crystallization trials and crystal optimization BacNavs pores were expressed and purified as described above. Seleno-methionine (SeMet) labelled proteins (CavSp1, NavAb1, NavAe1_Sp1CTD_p) were expressed following the metabolic inhibition protocol^56^. Due to general lower expression levels of SeMet labelled proteins, each purification required 6 L cultures and yielded around 20 µL at 7.5-10 mg mL^-1^. Purification was carried out as described above with the addition of 1mM tris(2-carbossietil)phosphine (TCEP) in each purification buffer.

##### CavSp1p

CavSp1p purified in DM as described above was concentrated to 13 mg mL^-1^ and reconstituted in bicelles^57^ prior crystallization to a final bicelle concentration of 8%. Native crystals grew in 25% PEG4000, 200 mM MgCl_2_, 100 mM MES, pH 6.5. To determine the three-dimensional structure, crystals of SeMet-labelled CavSp1p were obtained from seeding with native crystals in the same crystallization condition.

##### NavAb1p

SeMet labelled NavAb1 purified in DM was concentrated to 13 mg mL^-1^ and crystallized in were 3% PEG300, 0.75M ammonium sulfate.

##### NavAe1_Sp1CTD_p:SAT09 complex

Purified NavAe1_Sp1CTD_p was incubated with a 1:1.5 molar excess of SAT09 Fab and the complex was isolated by size exclusion chromatography using a Superdex200 (200mM NaCl, 0.3mM DDM, 20mM Hepes pH 8.0) and concentrated to 22-23 mg mL^-1^. Crystals grew at 20°C in 12-14% PEG4000, 0.1M NaCl, 0.1M MgCl_2_, 1 M Na acetate, pH 3.2-3.6. Crystals were harvested at 4°C by adding a cryoprotectant solution containing 17% PEG4000, 1mM DDM, 0.1M NaCl, 0.1M MgCl_2_, 0.1 M Na acetate, pH 3.6 and 5%-increasing steps of glycerol to a final concentration of 25%.

##### NavAe1_Sp1CTD_p:ANT05 complex

Purified NavAe1_Sp1CTD_p was incubated with a 1:1.5 molar excess of ANT05 Fab and the complex was isolated by size exclusion chromatography on a Superdex200 (200 mM NaCl, 0.3 mM DDM, 20mM Hepes, pH 8) and concentrated to 36 mg mL^-1^. Crystals grew at 20°C in 13% PEG4000, 2% propanol, 0.1M LiS0_4_, 0.1 M 2-[(2-amino-2-oxoethyl)-(carboxymethyl)amino]acetic acid (ADA) pH 6.5. Crystals were harvested at 4°C by adding a cryoprotectant solution containing 13% PEG4000, 2% propanol, 1mM DDM, 0.1M LiS0_4_, 0.1 M ADA pH 6.5, and 5%-increasing steps of glycerol to a final concentration of 25%.

##### NavAe1_Sp1CTD_p

Purified NavAe1_Sp1CTD_p was concentrated to 13-13.5 mg mL^-1^ and crystallized in 22% PEG3350, 0.3 M KI, 8 mM sarcosine. Crystals were harvested in 30% PEG3350, 0.3 M KI, 8 mM sarcosine and 1mM Fos-choline 12 (FC-12), Anatrace). The FC-12 was added in order to remove the skin covering each crystal, leading to higher diffraction. Seleno-methionine containing crystals diffracted at higher resolution than the native ones and were used for structure determination.

#### Structure determination

##### CavSp1p and NavAb1p

Data from both native and Se-Met derived crystals were collected at the Advanced Light Source Beamline 8.3.1, Lawrence Berkeley National Laboratory. 2- and 3-wavelength MAD datasets were collected for the Se-Met crystals using the Selenium peak and inflection, or peak, inflection, and remote wavelengths for CavSp1p and NavAb1p, respectively. Diffraction images were integrated using iMOSFLM 1.0.5^58^, and scaled with SCALA (3.3.20)^59^. Initial experimental phases were determined using SHELXE^60^. An initial model was obtained by manual placing secondary structure elements and improved with iterative rounds of manual rebuilding with COOT^61^ and refinement with Refmac (5.6.0117)^62^.

##### NavAe1_Sp1CTD_p:SAT09 complex and NavAe1_Sp1CTD_p:ANT05 complex

Diffraction data for NavAe1_Sp1CTD_p:SAT09 and NavAe1_Sp1CTD_p:ANT05 complexes were collected at the Advanced Photon Source Beamlines 23ID-B and 24ID-E, respectively. Data were processed with XDS^63^, scaled and merged with Aimless^64^. Structures were solved by molecular replacement using the Fab fragment (PDBID: 4XGZ and its stable H12^65^variant for the SAT09 and ANT05 complexes, respectively) as search model. Several cycles of manual rebuilding, using COOT^61^, and refinement using REFMAC5 (5.6.0117)^62^and PHENIX^66^ were carried out to improve the electron density map. For AeSpCTD:ANT05 complex, secondary structure and Ramachandran restraints together with four-fold NCS restraints and no side-chains were used to refine the final model.

##### NavAe1_Sp1CTD_ SeMet

Three MAD datasets corresponding to the peak, inflection and remote Selenium wavelengths were collected at the Advanced Photon Source Beamlines 23ID-B. Data were processed with XDS. Unmerged data were submitted to the automated structure solution pipeline CRANK2^67^ from the CCP4i2 interface to determine initial experimental phases. An initial model was obtained by manual placing secondary structure elements and improved with iterative rounds of manual rebuilding with COOT and refinement with Refmac (5.6.0117)^62^. The full model was then refined to convergence in CNS^68,69^ and PHENIX^66^ using secondary structure and Ramachandran restraints together with four-fold NCS restraints and a higher resolution reference model (PDBID:5HK7).

---

**Fig. S1.**
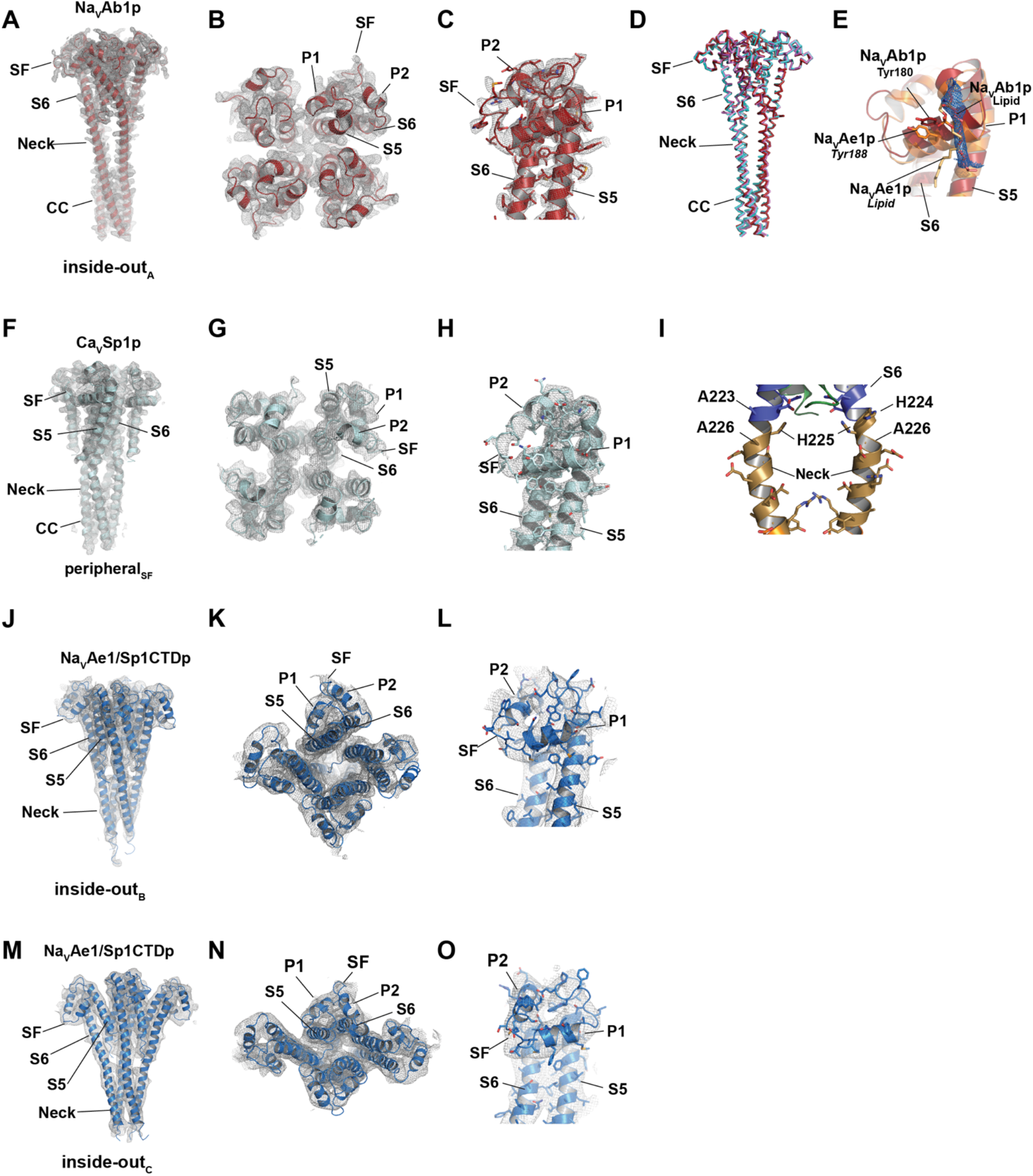
BacNav PDs exemplar electron density and structural details. **A-C,** 2Fo-Fc Electron density (1σ) for NavAb1p (firebrick) **A,** side view, **B,** extracellular view, and **C,** Single subunit. **D,** Superposition of NavAb1p structures determined in detergent (firebrick) and bicelles (cyan). **E,** Comparison of lipid bound to the P1 helix in NavAb1p (firebrick) and NavAe1p (PDB 5HK7)^1^. This lipid is modeled as phosphoenthanolamine and the 2Fo-Fo density (1.0σ, marine mesh) shows a well-defined acyl chain sitting between the P1 and S6 helices of NavAb1p. **F-H,** 2Fo-Fc Electron density (1σ) for CavSp1p (cyan) **F,** side view, **G,** extracellular view, and **H** Single subunit. **I,** Close up of the CavSp1p S6 (marine)-Neck (olive) junction showing two subunits. Select residues are indicated. **J-O,** 2Fo-Fc Electron density (1σ) for NavAe1/Sp1ctdp (marine). **J-L,** inside-outb tetramer **J,** side view, **K,** extracellular view, and **L,** Single subunit. **M-O,** inside-outc tetramer **M,** side view, **N,** extracellular view, and **O,** Single subunit.

**Fig. S2.**
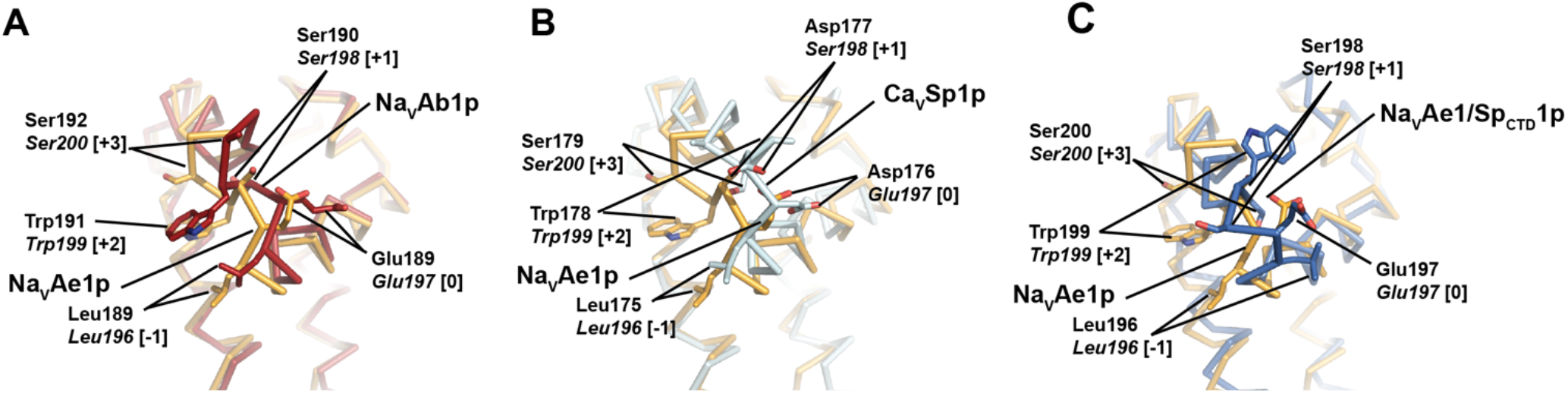
Structure comparison of selectivity filters from canonical and non-canonical quaternary assemblies. **A,** NavAb1p (firebrick), **B,** CavSp1p (cyan), and **C,** NavAe1/Sp1ctdp inside-outc form(marine) compared with the NavAe1p canonical structure (orange) (PDB:5HK7)^1^.

**Fig. S3.**
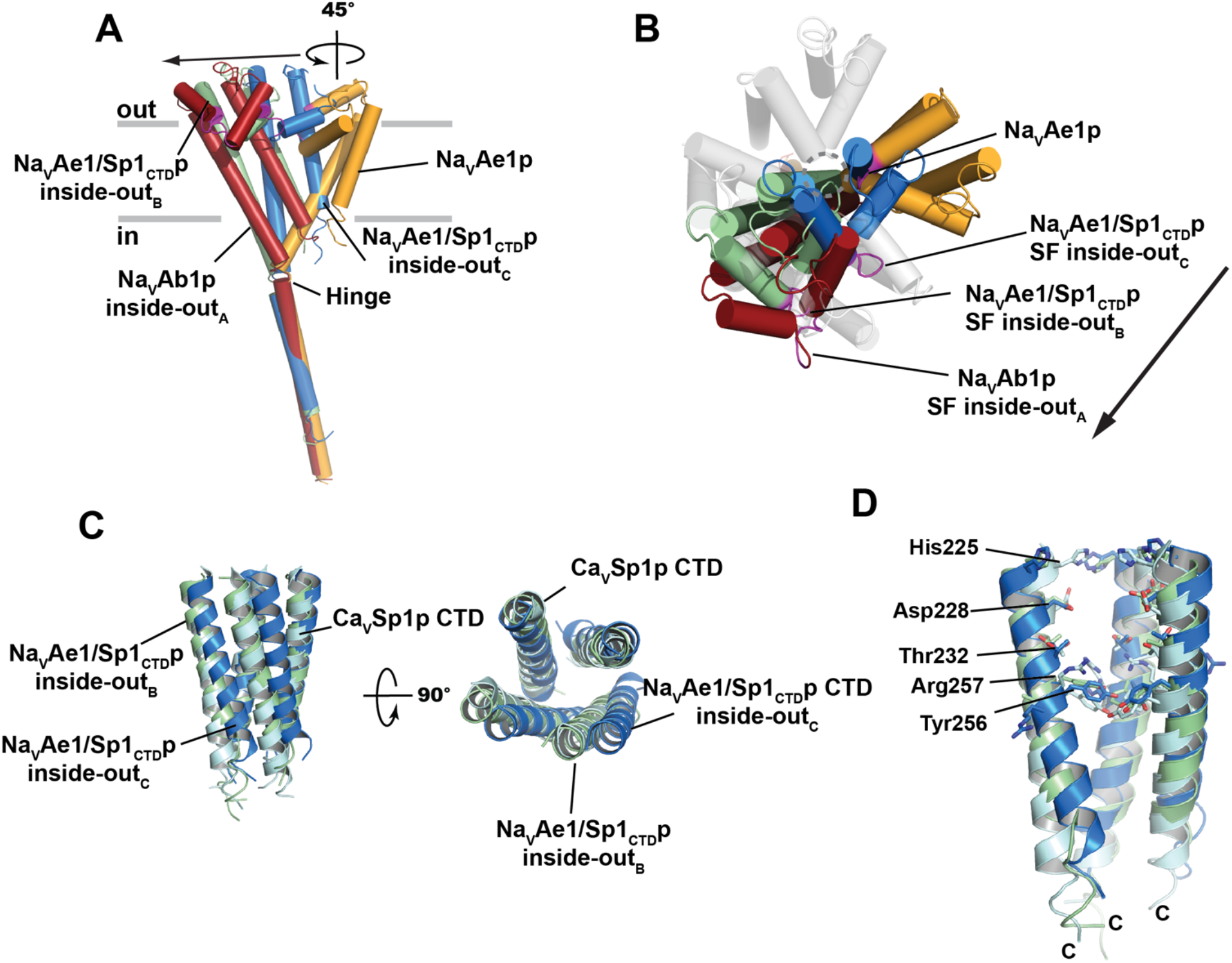
Structure comparison of inside-out and canonical quaternary assemblies. **A,** Superposition showing the rigid body movements that connect NavAe1p (orange), NavAb1p, inside-outa (firebrick), NavAe1/Sp1ctdp inside-outb (pale green), and NavAe1/Sp1ctdp inside-outc (marine) conformations. C-tails of each monomer are superposed. The selectivity filter of each monomer is magenta. All inside-out forms are related to NavAe1p by a 45° rotation round the hinge followed by varied degrees of translation indicated by the arrow. **B,** Extracellular view of ‘A’. Location of central ion conducting pore is indicated by the open circle. NavAe1p tetramer is shown with one orange and three white subunits. Arrow shows the relationships among the inside-out forms**. C,** Superposition of the CTDs from NavAe1/Sp1ctdp inside-outb (pale green) and inside-outc (marine) conformations with CavSp1p (cyan). **D,** Details of the central core from ‘C’.

**Fig. S4.**
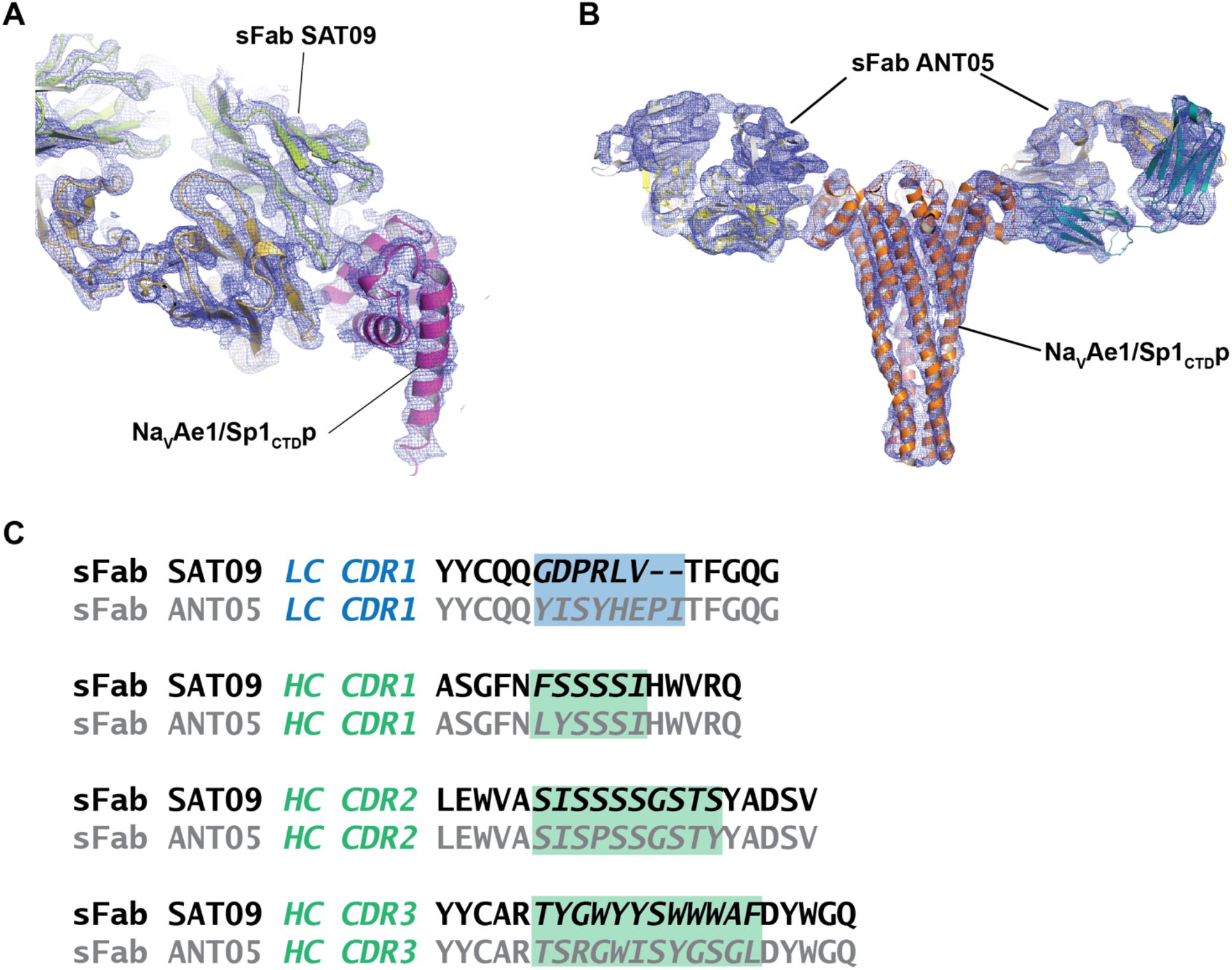
sFab:NavAe1/Sp1ctdp complexes. **A,** 2Fo-Fc (2σ) electron density for sFab SAT09:NavAe1/Sp1ctdp. sFab SAT09 light (limon) and heavy (yellow orange) chains and NavAe1/Sp1ctdp (magenta) are indicated. **B,** 2Fo-Fc (1σ) electron density for sFab ANT05:NavAe1/Sp1ctdp. sFab ANT05 light (aquamarine) and heavy (yellow) chains and NavAe1/Sp1ctdp (orange) are indicated. **C,** Sequence comparisons of the light chain (LC) and heavy chain (HC) CDRs (blue and green, respectively) for sFabs SAT09 (black) and ANT05 (grey). CDR sequences are shown in italics.

**Fig. S5.**
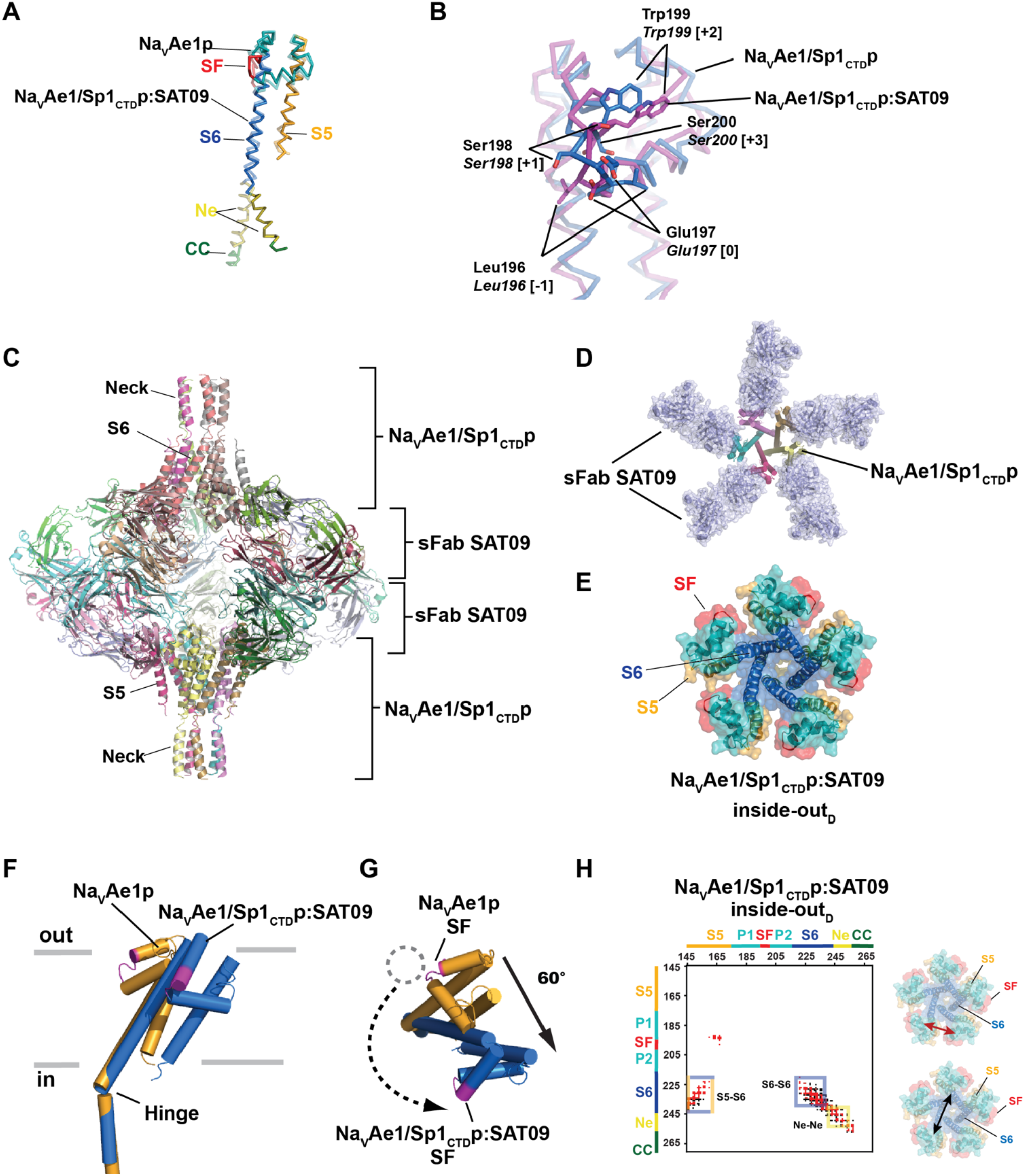
Structure of the sFabSAT09:NavAe1/Sp1ctdp complex. **PD. A,** Superposition of NavAe1/Sp1ctdp from the sFab SAT09:NavAe1/Sp1ctdp complex and NavAe1p from the canonical structure (PDB:5HK7)^1^. Channel elements are colored as follows, S5 (bright orange), SF (red), P1 and P2 helices (teal), S6 (marine), neck (olive) coiled-coil (forest).**B,** Superposition of NavAe1/Sp1ctdp (marine) and NavAe1/Sp1ctdp from the SAT09 complex (magenta). **C**, sFabSAT09:NavAe1/Sp1CTDp complex asymmetric unit. **D**, Extracellular view of a sFabSAT09:NavAe1/Sp1CTDp pentameric complex (top). **E**, Extracellular view of a NavAe1/Sp1CTDp inside-out_D_. Channel elements are colored as in ‘A’. **F,** Superposition showing the rigid body movements that connect conformations of NavAe1p (orange) and NavAe1/Sp1ctdp from the sFabSAT09:NavAe1/Sp1ctdp complex (marine). C-tails of each monomer are superposed. The selectivity filter of each monomer is magenta. Hinge is indicated. **G,** Extracellular view of ‘F’. Location of central ion conducting pore in NavAe1p is indicated by the open circle. Arrows show the NavAe1p-NavAe1/Sp1ctdp relationship**. H**, sFabSAT09:NavAe1/Sp1ctdp complex contact map. C_α_-C_α_ distances for (black) diagonal subunits at 20 Å and (red) neighboring subunits at 12 Å. Channel structural elements are indicated. Extracellular views of the PDs having channel elements colored as in ‘A’ are shown. Arrows indicate the diagonal (black) and neighbor (red) distance relations of the contact plots.

**Fig. S6.**
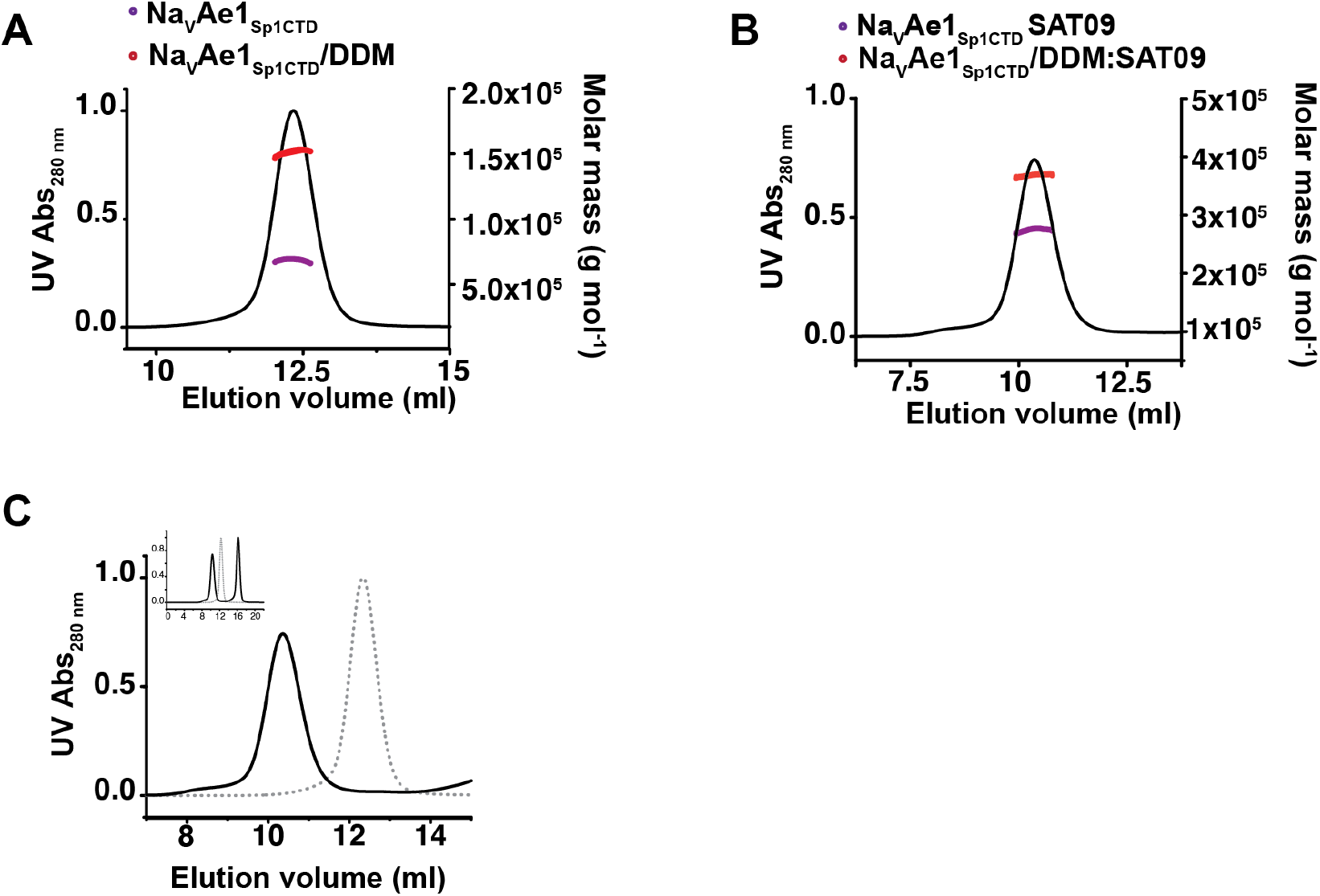
SEC-MALS analysis of NavAe1_Sp1Ctd_ and SAT09:NavAe1_Sp1CTD_. **A,** SEC-MALS chromatograms of 15 µM NavAe1_Sp1CTD_ purified in DDM. **B,** SEC-MALS chromatograms of 15 µM NavAe1_Sp1CTD_ in complex with 2.5-fold excess of sFab SAT09. The red and purple lines represents respectively the total molar mass and protein molar mass fitting results. **C,** Superimposition of NavAe1_Sp1CTD_ and SAT09:NavAe1_Sp1CTD_ SEC-MALS chromatograms from ‘A’ and ‘B’.

**Fig. S7.**
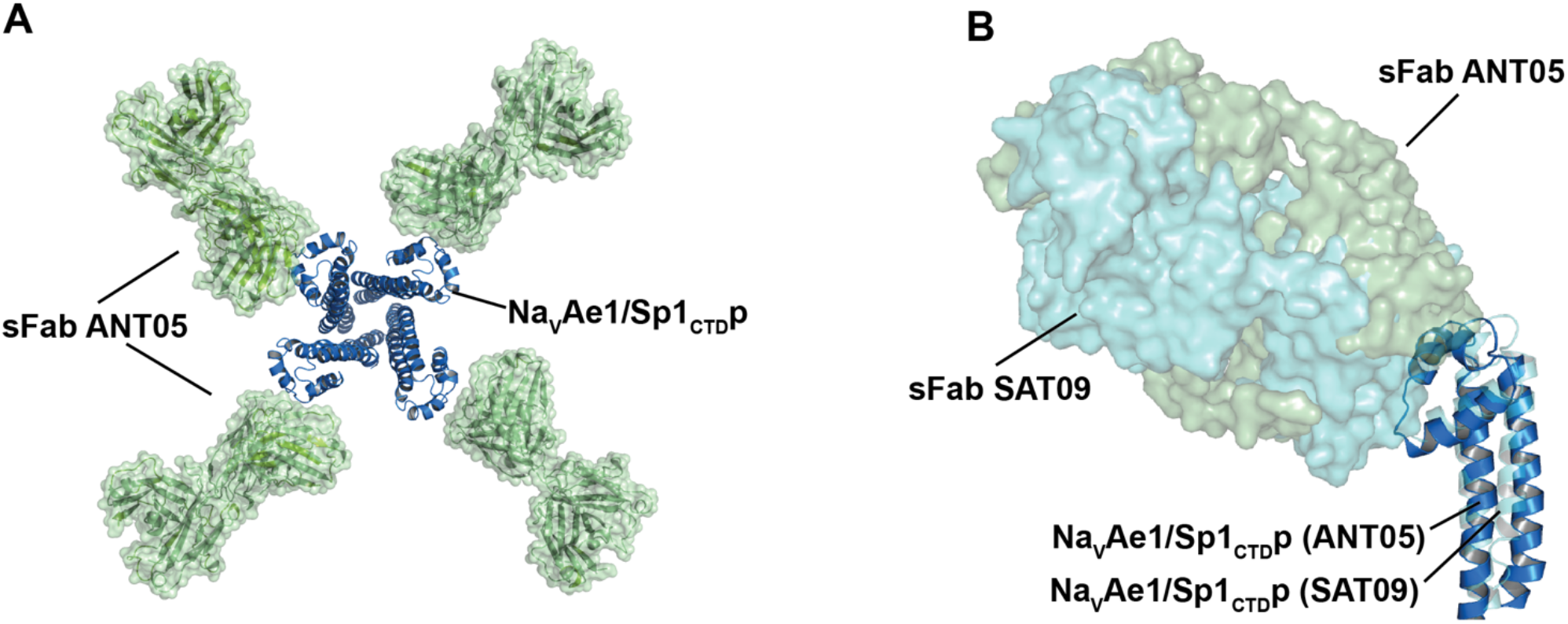
Structure of the sFabANT05:NavAe1/Sp1CTDp complex. **A**, Extracellular view of the sFabANT05:NavAe1/Sp1CTDp complex. sFabANT05 (green) is shown in cartoon and surface rendering. NavAe1/Sp1CTDp (marine) is shown in cartoon rendering. **B,** Comparison of the binding modes of sFabANT05 (green) and sFabSAT09 (cyan) to NavAe1/Sp1CTDp. NavAe1/Sp1CTDp from the sFabANT05 complex is marine. NavAe1/Sp1CTDp from the sFabSAT09 complex is cyan.

**Fig. S8.**
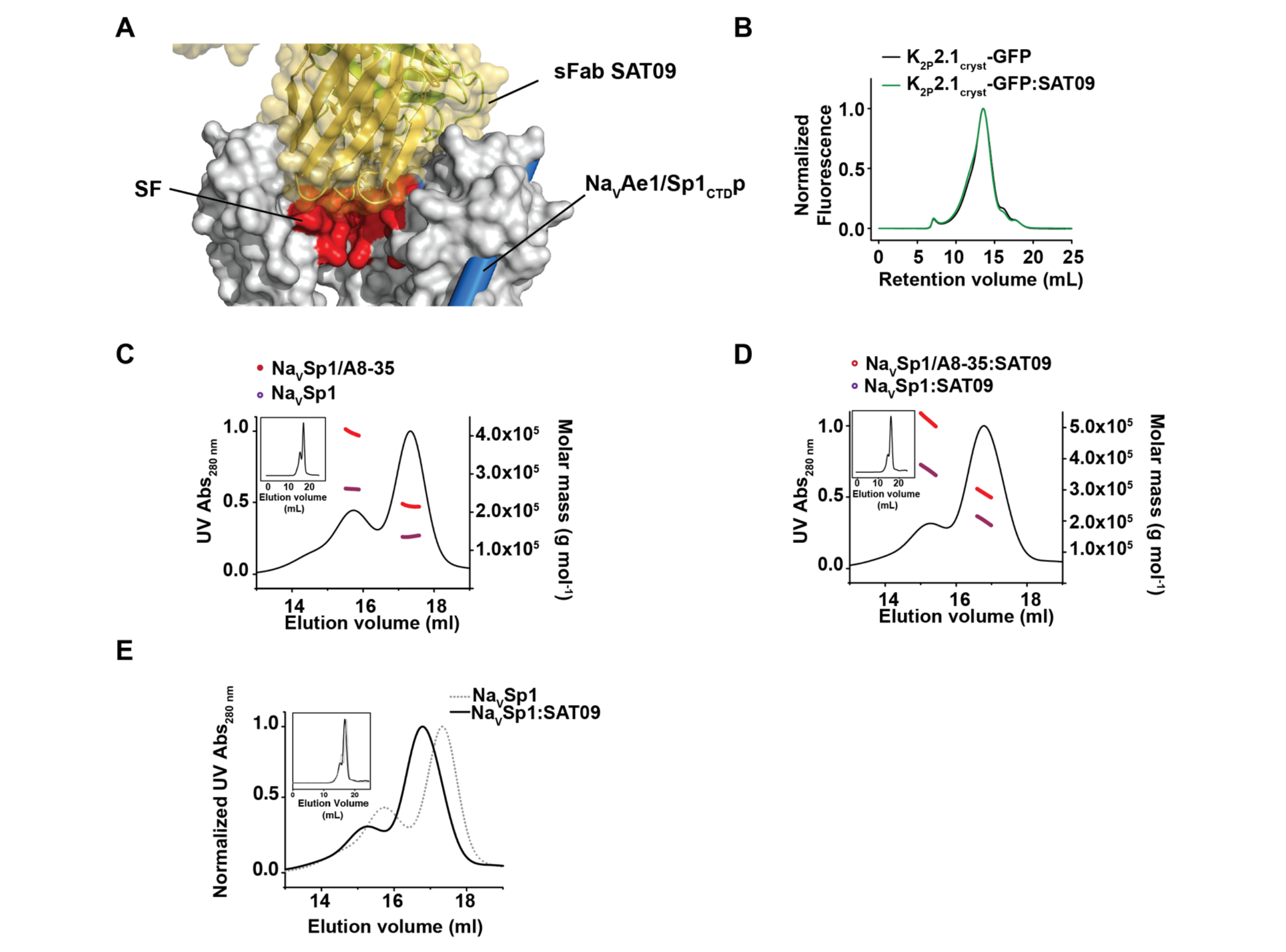
Characterization of sFab SAT09 binding. **A,** Superposition of the sFabSAT09 complex (solid yellow, green SAT09 and marine cylinders NavAe1/Sp1CTDp on the NavAe1p (space filling, white) canonical structure (PDB:5HK7)^1^ showing three of the four subunits. Selectivity filter (SF) region of NavAe1p is colored red. **B,** Superose 6 10/300 FSEC profiles of K_2P_2.1(TREK-1)_cryst_-GFP alone (black) and with SAT09 (green). DDM-solubilized fraction from K_2P_2.1(TREK-1)_cryst_-GFP expressing *Pichia p.* cells (100 µl) was incubated with 1 nmol of sFab SAT09. **C,** SEC-MALS chromatograms of 9 µM NavSp1 reconstituted in amphipol A8-35. **D,** SEC-MALS chromatogram of 9 µM NavSp1-SAT09 complex reconstituted in amphipol A8-35 and taken after purification of the complex on Superose 6. **E,** Superimposition of NavSp1 (dashed line) and SAT09:NavSp1 (black) SEC-MALS chromatograms from ‘C’ and ‘D’.

**Fig. S9.**
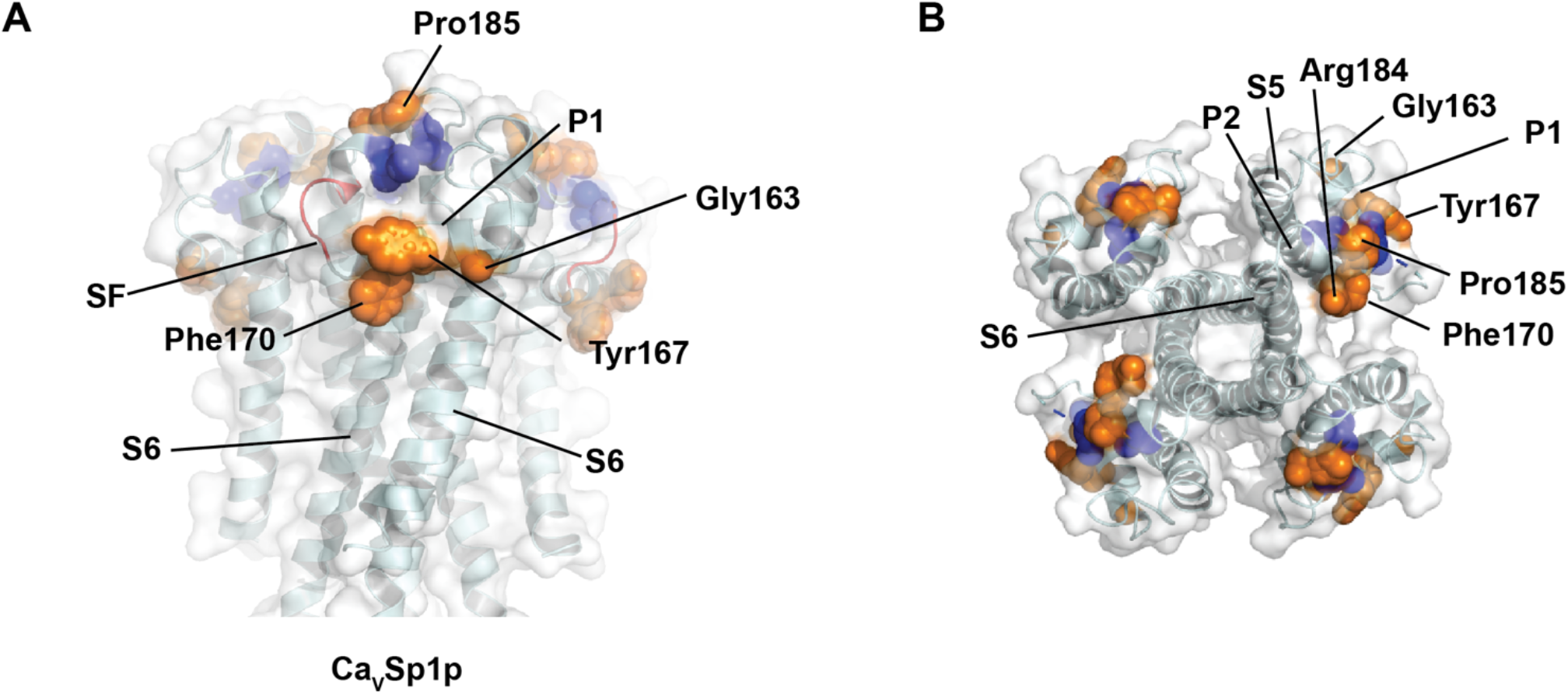
EPR mobility changes. **A,** Side and **B,** Extracellular views of the CavSp1p structure showing residues having changed mobility relative to the full length channel. Increased (orange), decreased (blue). Selectivity filter is colored red. EPR data are from^2^.

**Table S1.** Crystallographic data collection and refinement statistics

|  |  |  |  |  |  |  |
| --- | --- | --- | --- | --- | --- | --- |
|  | NavAb1p<br>detergent (DM)<br>(PDB: 7PGG) | NavAb1p<br>(bicelles)<br>(PDB: 7PGI) | CavSp1p (bicelles)<br>(PDB: 7PGF) | NavAe1/Sp1<sub>CTDP</sub><br>(DDM)<br>(PDB: 7PGH) | NavAe1/Sp1<sub>CTDP</sub><br>:SAT09 complex<br>(PDB: 7PGP) | NavAe1/Sp1<sub>CTDP</sub><br>:ANT05 complex<br>(PDB:7PG8) |
| **Data Collection** |  |  |  |  |  |  |
| Space group | P4<sub>3</sub>2<sub>1</sub>2 | I2<sub>1</sub>2<sub>1</sub>2<sub>1</sub> | P6<sub>1</sub>22 | P2<sub>1</sub>2<sub>1</sub>2<sub>1</sub> | P4<sub>3</sub> | P4<sub>3</sub> |
| Cell dimensions a/b/c (Å) | 157.471/ 157.471/<br>106.394 | 178.179/ 191.80<br>/ 192.32 | 133.682/ 133.682/<br>130.744 | 123.465 /134.696/<br>155.545 | 200.865 / 200.865/<br>327.727 | 127.175/ 127.175/<br>445.206 |
| $\alpha/\beta/\gamma$ (°) | 90/ 90/ 90 | 90/ 90/ 90 | 90/ 90/ 90 | 90/90/90 | 90/ 90/ 90 | 90/ 90/ 90 |
| Resolution (Å) | 14.96 - 2.852<br>(2.953 - 2.852) | 19.98 - 3.64<br>(3.769 - 3.64) | 12.48 - 3.5 (3.622 -<br>3.5) | 14.99 - 4.2 (4.342 -<br>4.195) | 15 - 3.6 (3.727 - 3.6) | 122.3 - 4.46 (4.62 -<br>4.46) |
| Rmerge (%) | 0.169 | 0.157 | 0.156 | 0.065 | 0.166 | 0.031 |
| I / $\sigma$ I | 15.4 (1.0) | 5.2 (0.9) | 10.2 (0.6) | 11.6 (1.1) | 9.3 (0.5) | 12.8 (0.7) |
| CC(1/2) | 0.999 (0.473) | 0.995 (0.172) | 0.995 (0.126) | 0.999 (0.404) | 0.998 (0.130) | 0.997 (0.442) |
| Completeness (%) | 98.99 (99.87) | 98.41 (98.88) | 97.17 (100.00) | 96.53 (90.59) | 98.47 (99.34) | 98.31 (88.69) |
| Redundancy | 16.2 (16.6) | 4.2 (3.8) | 11.5 (11.9) | 12.9 (12.7) | 7.0 (6.8) | 2.0 (1.9) |
| Unique reflections | 31433 (3087) | 36662 (3616) | 8915 (879) | 18983 (1766) | 147489 (14754) | 42377 (3816) |
| Wilson B-factor | 95.36 | 122.90 | 134.30 | 267.01 | 156.76 | 254.31 |
| Wavelength (Å) | 1.115869 Å | 1.11587 Å | 1.11587 Å | 1.0000 Å | 1.0000 Å | 1.0000 Å |
| **Refinement** |  |  |  |  |  |  |
| R<sub>work</sub> / R<sub>free</sub> (%) | 21.5/23.8 | 28.7/30.4 | 20.9/24.4 | 32.8/34.30 | 20.8/23.4 | 36.4/39.8 |
| No. of chains in AU | 2 | 8 | 2 | 8 | 30 | 24 |
| No. of protein atoms | 2363 | 9291 | 2029 | 8413 | 42648 | 22623 |
| No. of ligand atoms | 21 | 1 | 1 | 415 | 1197 | 0 |
| No. of water atoms | 0 | 0 | 1 | 0 | 9 | 0 |
| RMSD bond lengths (Å) | 0.017 | 0.014 | 0.012 | 0.006 | 0.004 | 0.004 |
| RMSD angles (°) | 2.59 | 1.90 | 1.75 | 1.20 | 1.08 | 0.76 |
| Ramachandran<br>best/disallowed regions (%) | 85.1/2.1 | 97.7/0.1 | 70.2/6.2 | 82.8/1.6 | 94.1/0.2 | 87.9/0.3 |

15 August 2021
|  |  |  |  |  |  |  |  |  |
| --- | --- | --- | --- | --- | --- | --- | --- | --- |
|  | NavAe1<sub>Sp1CTDp</sub> DDM |  |  | NavAb1p, detergent (DM) |  |  | CavSp1p bicelles |  |
| Data Collection | inflection | peak | remote | inflection | peak | remote | peak | remote |
| Space group | P 21 21 21 | P 21 21 21 | P 21 21 21 | P 41 21 2 | P 41 21 2 | P 41 21 2 | P 61 2 2 | P 61 2 2 |
| Cell dimensions a/b/c (Å) | 125.43<br>/135.80/<br>156.13 | 125.12 /<br>135.38/<br>155.76 | 125.31/<br>135.64/<br>156.00 | 157.6260<br>157.6260<br>106.5250 | 157.2440<br>157.2440<br>106.0030 | 157.3780<br>157.3780<br>106.3530 | 133.0465 133.0465<br>128.1215 | 133.08 133.08<br>128.09 |
| $\alpha/\beta/\gamma$ (°) | 90/90/90 | 90/ 90/ 90 | 90/ 90/ 90 | 90/90/90 | 90/ 90/ 90 | 90/ 90/ 90 | 90/ 90/ 90 | 90/ 90/ 90 |
| Resolution (Å) | 48.89 - 4.51<br>(5.04 - 4.51) | 48.77 - 4.51<br>(5.04 - 4.51) | 48.85 - 4.51<br>(5.04- 4.51) | 39.41 - 3.28<br>(3.46 - 3.28) | 39.31 - 3.20<br>(3.37 - 3.20) | 39.34 - 3.20<br>(3.37 - 3.2) | 34.31 - 3.94 (4.40 - 3.94) | 34.31 - 3.85 (4.30 - 3.85) |
| Rmerge (%) | 0.044 (>1) | 0.041 (>1) | 0.043 (>1) | 0.260 (>1) | 0.182 (>1) | 0.239 (>1) | 0.349 (>1) | 0.401 (>1) |
| I / $\sigma$ I | 10.8 (0.6) | 11.6 (1.0) | 11.4 (0.8) | 14.2 (1.0) | 19.5 (2.2) | 15.7 (0.8) | 16.6 (1.1) | 15.2 (0.9) |
| CC(1/2) | 0.999 (0.286) | 1.0 (0.439) | 1.0 (0.370) | N/A | N/A | N/A | 1.00 (0.282) | 1.00 (0.228) |
| Completeness (%) | 99.40 (98.30) | 99.60 (98.80) | 99.20 (97.70) | 98.7 (92.0) | 99.9 (100.0) | 99.8 (99.6) | 98.3 (94.7) | 98.4 (95.1) |
| Redundancy | 6.6 (6.7) | 6.6 (6.8) | 6.6 (6.7) | 33.5 (32.2) | 34.1 (35.8) | 33.4 (31.7) | 64.6 (33.7) | 64.6 (33.6) |
| Unique reflections | 16246 (4499) | 16137 (4463) | 16169 (4454) | 20803 (2766) | 22495 (3216) | 22586 (3215) | 6214 (1630) | 6659 (1760) |
| Wilson B-factor | 351.6 | 294.9 | 328.4 | 166.3 | 128.5 | 166.1 | 145.7 | 145.2 |
| Wavelength (Å) | 0.97698 | 0.97953 | 0.95368 | 0.97960 | 0.97940 | 0.96110 | 0.979764 | 0.957038 |
| Refinement |  |  |  |  |  |  |  |  |

**Table S2.**
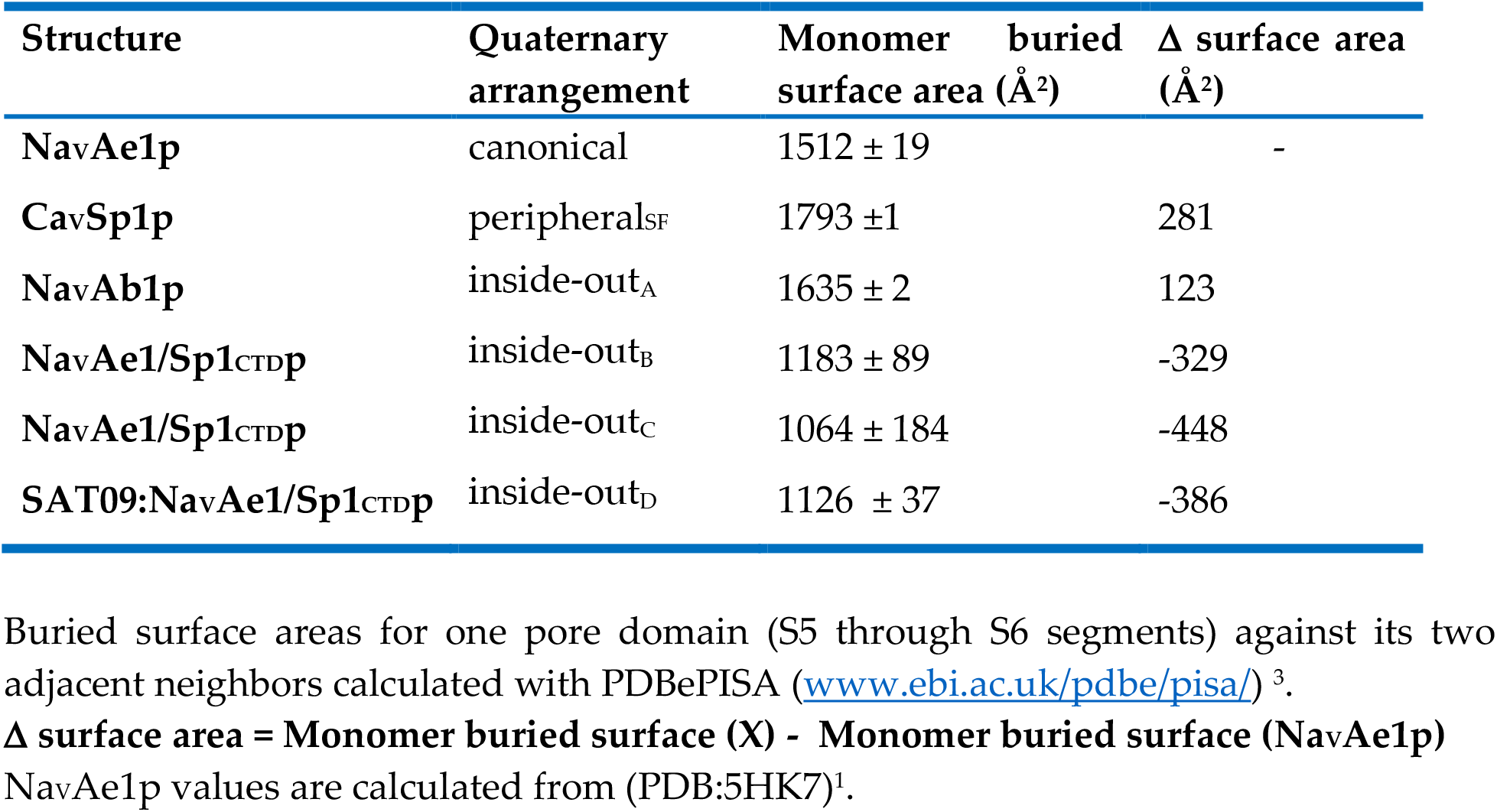
Pore domain monomer buried surface area

**Table S3.**
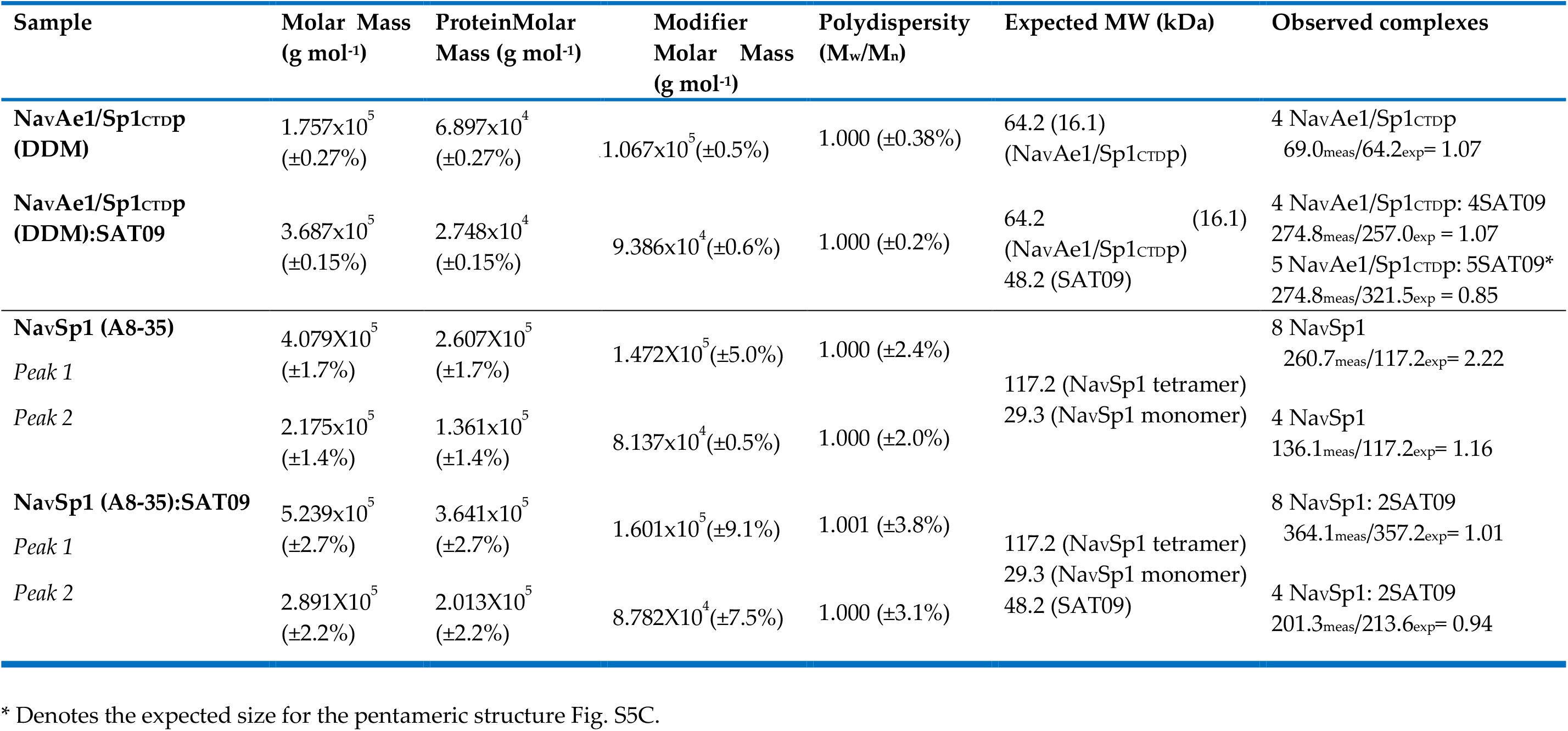
MALS data

